# Polytope: High-resolution epitope barcoding for *in vivo* spatial fate-mapping

**DOI:** 10.1101/2024.11.20.624484

**Authors:** Daniel Postrach, Colin E. J. Pritchard, Larissa Frank, Tom van Leeuwen, Hendrik A. Messal, Paul Krimpenfort, Jacco van Rheenen, Hans-Reimer Rodewald

## Abstract

Tracing the fate of individual cells and their progeny remains a challenging task. While imaging-based fate-mapping provides spatial information, it generally lacks complexity due to limited label diversity, resulting in diminished capture of comprehensive lineages and fate maps. Here, we introduce ‘Polytope’, an epitope barcoding system capable of generating up to 512 unique color codes. Comprising nine epitope tag cassettes flanked by loxP sites, Polytope allows random excision via Cre recombinase, creating unique color codes detectable through multiplexed imaging. Using an engineered Polytope mouse, we traced the fate of hundreds of clones across tissues *in vivo*, from embryonic development until adulthood. Together, Polytope enables high-resolution imaging-based fate-mapping through endogenous epitope barcoding, comprehensively capturing the spatial organization of complex clonal dynamics *in situ*.

**One-Sentence Summary:** Polytope is an imaging-based barcoding system to trace lineages at clonal and spatial resolution in living organisms using a diverse set of color-coded markers.

---

The ability to trace the developmental trajectories and spatial organization of cells within complex organisms is central to understanding tissue dynamics in health and disease (*1*, *2*). Fate-mapping technologies have become invaluable for studying stem cell differentiation and lineage contributions over time (*3–5*). Classical approaches, such as single-fluorescent reporter systems, provide insights into precursor-progeny relationships at the population level but lack the resolution required to trace the output of individual progenitors (*6*, *7*). Advanced imaging-based fate-mapping systems, such as the Confetti mouse model, introduced four-color fluorescent reporters, allowing *in vivo* visualization of individual cells and their progeny (*8*, *9*). However, these systems remain limited by the number of distinguishable colors, resulting in insufficient clonal resolution in most tissues.

Recent advances in cellular barcoding have improved fate-mapping by introducing unique molecular tags into individual cells, enabling their descendants to be tracked at exceedingly high clonal resolution (*10–19*). These methods rely on DNA barcodes, which are stably integrated into the genome and passed on to daughter cells, allowing for retrospective identification of clonal relationships. Despite their high precision in distinguishing cellular clones, they generally require sample destruction, such as during sequencing, which results in the loss of spatial information. Meanwhile, the rise of spatial biology technologies, such as multiplexed imaging, has enabled *in situ* mapping of tissues at single-cell resolution (*20*, *21*). Although these spatial approaches provide detailed snapshots of tissue organization, they typically do not capture lineage relationships or the clonal history of the cells. Bridging this gap requires new approaches that integrate cellular barcoding with spatially resolved imaging techniques, thereby linking the spatial context with clonal fate maps.

Here, we introduce ‘Polytope’, a spatial fate-mapping system based on *in vivo* epitope barcoding. Polytope leverages multiplexed imaging to resolve the spatial dynamics of clones *in situ* by generating up to 512 unique color codes in individual cells through Cre/loxP recombination. Polytope barcodes, introduced into stem and progenitor cells, are stably inherited to daughter cells, allowing for retrospective analysis of lineage relationships and clonal kinetics at high resolution. This technology enables the overlay of fate maps with spatial data obtained from multiplexed imaging, integrating clonal information with cell type annotations and cell states. Thereby, Polytope barcoding captures the fate, location and identity of cells on tissue sections, providing a comprehensive understanding of tissue dynamics during development and homeostasis *in vivo*.

## Random recombination of nine epitope tag cassettes generates 512 color codes

Polytope comprises nine epitope tag cassettes – FLAG, HA, V5, T7, VSVg, AU1, Myc, S-tag, and HSV – each flanked by loxP sites with a consistent spacing of 179 base pairs (Fig. 1A, B), similar to the core design of Polylox (*10*). LoxP sites are uniformly oriented, permitting excision of cassettes by Cre recombinase. Each cassette begins with an alpha-helix-forming EAAAK linker sequence, followed by 2 to 3 repeats of the specific epitope tag sequence and another EAAAK linker sequence (Fig. 1B). The entire construct, including the loxP sites, is expressed in the nucleus as an 84 kDa histone 2B (H2B) fusion protein (Fig. 1A). We also tested other linker designs, including the flexible GGGGS linker (*22*, *23*), which, however, resulted in protein degradation. We conclude that the rigid EAAAK linker stabilizes the peptide which is key for the expression and hence utility of Polytope.

**Fig. 1:**
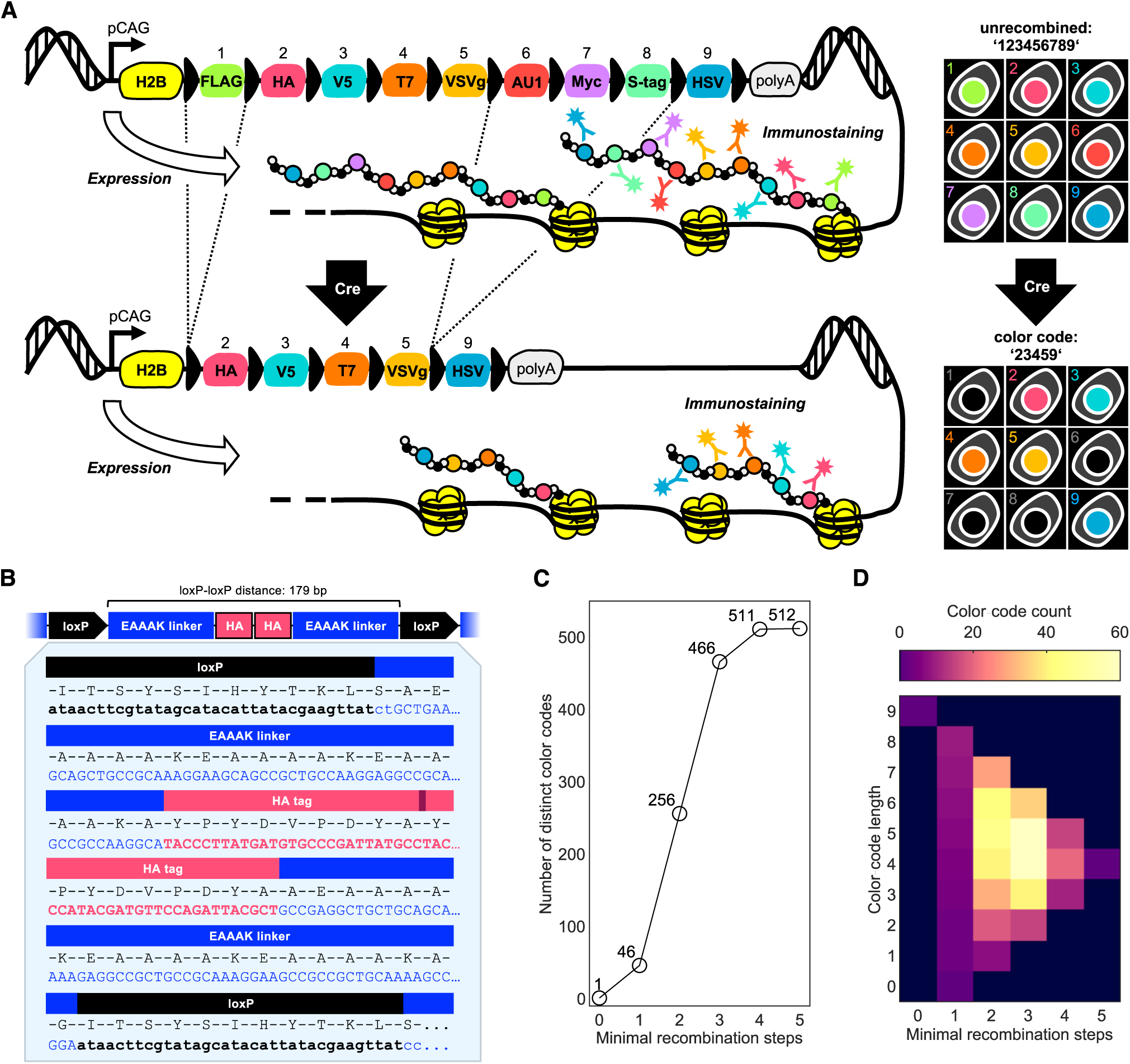
The principle and theory of Polytope epitope barcoding. (**A**) The Polytope peptide consisting of nine distinct epitope tags is expressed in the nucleus as an H2B fusion peptide. Color codes are generated through random loss of epitope cassettes upon Cre/loxP recombination and are detected by multiplexed imaging. (**B**) Exemplary design of the epitope tag cassettes with a consistent spacing of 179bp between loxP sites. 2-3 epitope tag sequences are flanked by rigid EAAAK linker sequences. (**C**) Up to 512 theoretically possible color codes are generated within a maximum of five consecutive recombination steps. (**D**) Cassette length and minimal recombination step distributions for all 512 color codes, indicating that the majority of all possible color codes consist of 3-6 epitopes and are generated within 2-3 recombination steps.

Based on this design, random excision of epitope tag cassettes from the locus by transient Cre activity (*10*, *24*) would generate distinct sets of epitope tag expressions, or color codes, which can be detected by multiplexed immunostaining (Fig. 1A). We define the color code according to the position of the epitope tag within the unrecombined locus (FLAG = 1, HA = 2, …). For instance, starting with the unrecombined sequence ‘123456789’, two successive recombination events might lead to the loss of the FLAG (’1’) and the AU1/Myc/S-tag (’678’) cassettes, generating the color code ‘23459’ (this theoretical color code is depicted in Fig. 1A). In up to five recombination steps, a maximum of 512 color codes can be generated (Fig. 1C, Table S1). This includes the possibility of fully recombining all available loxP sites, resulting in the loss of all nine epitope tags, which we define as the color code ‘0’. The majority of all possible color codes consists of 3-6 epitope tags and is generated within 2-3 recombination steps (Fig. 1D), suggesting that efficient yet diverse color code generation is experimentally achievable.

## Polytope color code induction and detection *in vitro*

We initially evaluated the expression and functionality of the Polytope system in human embryonic kidney (HEK) cells by targeting the open harbor locus AAVS1 (*25*) using CRISPR/Cas9 to create an *AAVS1^Polytope/+^* knock-in cell line. Polytope color code detection was achieved by immunostaining all nine epitope tags using the IBEX iterative immunolabeling and chemical bleaching method (*26*, *27*) in sets of three tag antibodies (Fig. 2A; Fig. S1A; Table S2). Indeed, in the *AAVS1^Polytope/+^* cell line, all nine epitopes are co-expressed in the nucleus (Fig. 2B). To induce recombination, *AAVS1^Polytope/+^* cells were treated with cell-permeable tat-Cre protein (*28*). Subsequent analysis by IBEX multiplexed imaging revealed diverse, visually distinct color codes in the fluorescent microscopy images (Fig. S1B, movie 1). An overview of the barcoded cells in culture, as well as individual regions, i.e. clones, at one week after color code induction are depicted in Fig. 2C. Cells in region 1 are unrecombined, while those in regions 2-5 show distinct color codes, including ‘156789’, ‘15’, ‘123489’, or ‘0’ (Fig. 2C). By applying nuclear segmentation and image analysis for each cell in the entire culture, we measured mean fluorescent intensity (MFI) values of all nine epitope tag channels (Fig. 2D, S1C, D). Histograms reveal dose-dependent recombination evident by loss of epitope tags, compared to the uniform presence of all nine epitope tags in untreated cells (Fig. 2D). We next applied automated gating based on the bimodal histograms (red line in Fig. 2D) to classify cells as positive or negative for each epitope tag, and hence assigned individual color codes. Using this approach, we analyzed two independent samples (either treated with 100 or 200 U/mL tat-Cre) and detected a total of 431 unique color codes in the combined samples, representing 84% of the theoretical maximum (Fig. 2E). The absolute number of distinct color codes was similar for both tat-Cre doses (353 and 382 for 100 and 200 units, respectively) despite of different recombination rates (Fig. 2D). The frequencies of observed color codes spanned five orders of magnitude (Fig. 2F), with color codes such as ‘123456789’, ‘0’, ‘1’, and ‘9’ being most abundant, and color codes such as ‘3579’ or ‘2468’ being rare, indicating that color codes are generated at unequal rates. This variability can be attributed to differences in the number of recombination steps required to generate each color code, as previously described in Polylox barcoding (*10*). In general, color codes generated in fewer steps tend to be more abundant than those requiring a higher number of recombination steps. Indeed, a comparison of experimentally derived color code frequencies with the number of required recombination steps revealed a negative correlation (Fig. 2G). However, since color-coded clones are spatially distributed, filtering for generation probability may not be required to resolve individual clones. To demonstrate this, we performed simulations by randomly distributing experimentally derived color codes on a 2D grid according to their detected frequencies (Fig. 2H), followed by counting cases in which neighboring cells shared the same color code. Iterating this process one hundred times, we found that only 0.27% of cells shared a color code with a neighbor (Fig. 2I), confirming that Polytope’s label diversity is sufficient to spatially distinguish clones. Color code detection strongly relies on robust and error-free epitope immunostaining and multiplexed imaging. To test whether each color code matches its underlying DNA barcode, we generated single cell-derived clonal lines from tat-Cre-treated *AAVS1^Polytope/+^* HEK cells (Fig. S2A). DNA barcodes were analyzed by PCR and subsequent Sanger sequencing (Fig. S2B), which was correlated with color codes derived from IBEX multiplexed imaging (Fig. S2C). In all instances, DNA and epitope barcodes were found to match (Fig. S2C). In summary, we detected a large number of distinct color codes, setting the stage to explore Polytope for spatial fate-mapping experiments *in vivo*.

**Fig. 2:**
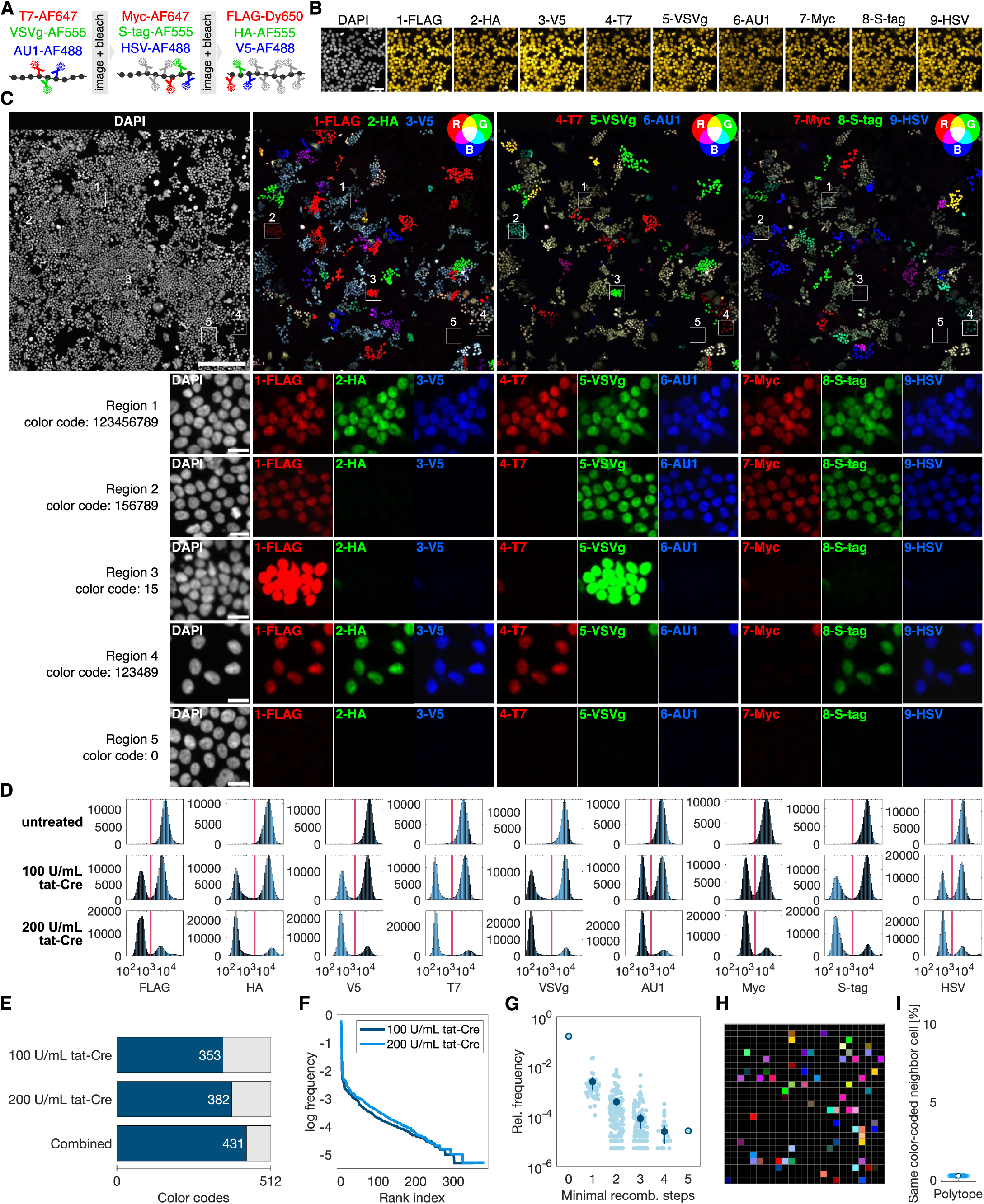
Polytope color coding *in vitro*. (**A**) Principle of the immunostaining-bleach cycles using the IBEX protocol for multiplexed imaging of all nine epitope tags. (**B**) IBEX multiplexed imaging of all nine epitope tags on fixed *AAVS1^Polytope/+^*HEK293 cells. Scale bar: 50 µm. (**C**) Tat-Cre treated *AAVS1^Polytope/+^*HEK293 cells and subsequent multiplexed imaging depicted in triplet overlays (RGB color system). Distinct regions of interest depict clones with varying color codes. Scale bar: 250 µm (large panel), 20 µm (ROI panels). (**D**) Histogram of mean fluorescent intensity signals of all nine epitope tags derived from image analysis. Red line indicates threshold for classification into either positive or negative. (**E**) Number of distinct color codes detected compared to the theoretical maximum. (**F**) Relative frequency of color codes detected in 100 U/mL and 200 U/mL tat-Cre treated samples. (**G**) Relative frequency of detected color codes compared to the number of minimal recombination steps required to generate the color code. (**H**) Example of random color code distribution simulated for detected color codes (each color indicates a distinct color code). **(I)** Frequency of same color-coded neighboring cells in random distribution simulation.

## Generation of a *Rosa26^Polytope/+^* knock-in mouse for *in vivo* color code induction

We generated a Polytope knock-in mouse line by targeting the Rosa26 locus in mouse embryonic stem cells via homologous recombination for integration of the Polytope construct (Fig. 3A). Successful integration was demonstrated by Southern blot analysis of tail DNA (Fig. 3B). Immunostaining of cryosections from various organs, including brain, intestine, kidney, liver, lung, skin, spleen and thymus, revealed consistent nuclear expression of the Polytope peptide in all tested tissues, with no detectable signal in wild-type controls (Fig. 3C). Systematic histopathological evaluation confirmed normal development and organ function in heterozygous and homozygous Polytope mice (Fig. S3).

**Fig. 3:**
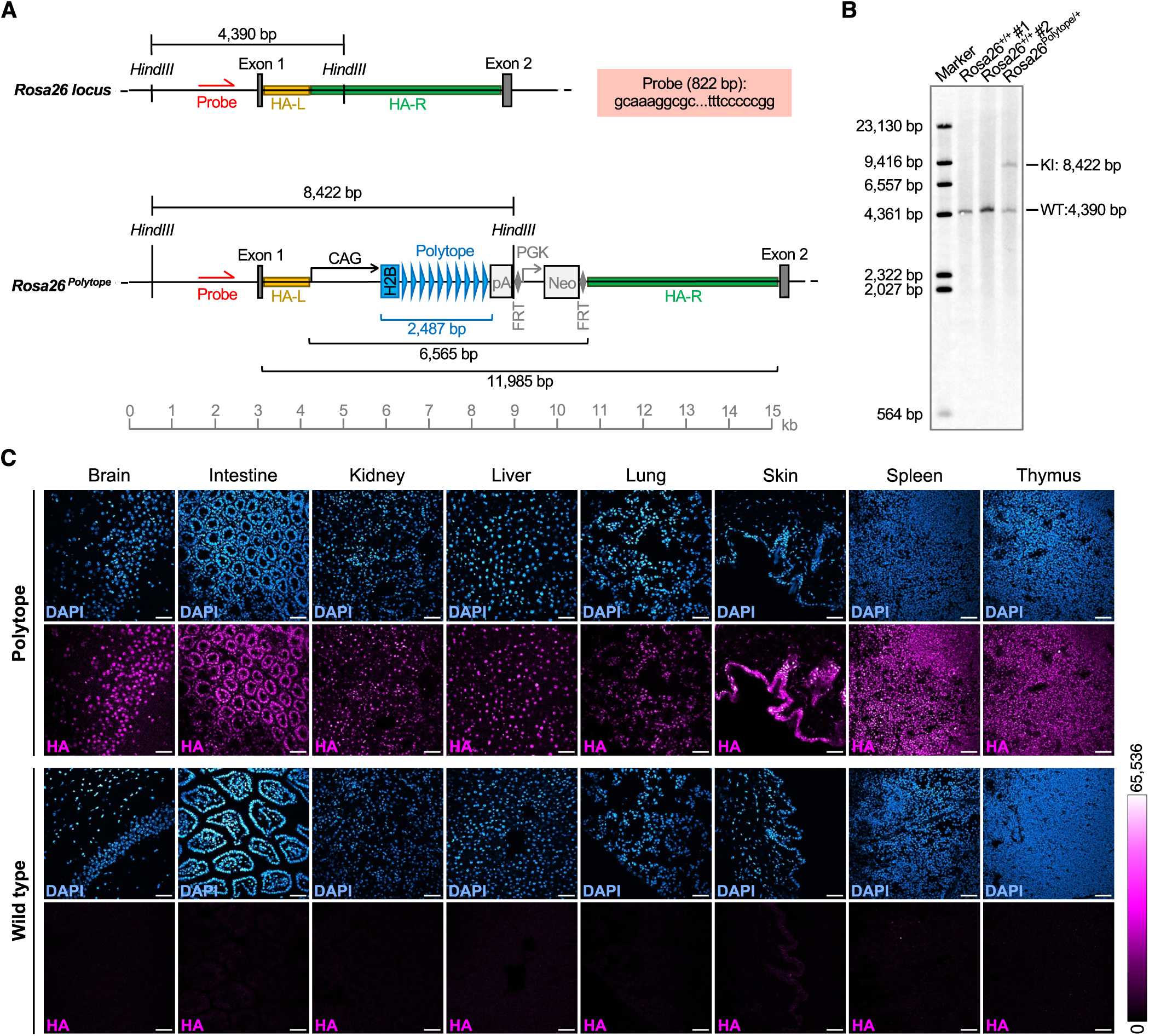
Generation of the *Rosa26^Polytope/+^* mouse. (**A**) Classical gene targeting of the Rosa26 locus in mouse embryonic stem cells by homologous recombination. The red box and arrow indicate the probe that was used in (B). HA-L/HA-R: homology arm left or right; CAG: CMV-early-enhancer/chicken-β-actin promoter; H2B: histone 2B cassette; FRT: FLP recombinase target; PGK: phosphoglycerate kinase 1 promoter; Neo: neomycin resistance gene. (**B**) Southern blot on genomic DNA of two wild-type (*Rosa26^+/+^*) mice and Polytope knock-in (*Rosa26^Polytope/+^*) mouse. Gene targeting was confirmed by the size of the restriction fragments corresponding to wild-type (4,390 bp) and knock-in (8,422 bp) using the probe indicated in (A). (**C**) Anti-HA tag immuno-staining and fluorescent microscopy on cryosection of various tissues derived from Polytope (*Rosa26^Polytope/+^*) and wild-type (*Rosa26^+/+^*) mice. Scale bar: 50 µm.

Next, we induced epitope barcoding using ubiquitously expressed inducible Cre in *Rosa26^CreERT2/Polytope^* mice by administering tamoxifen (TAM). To allow physiological clone formation and its visualization by color codes, mice were given a four-week chase period after TAM (Fig. 4A). Cryosectioning and immunostaining revealed distinct color-coded clones that had emerged in intestine (Fig. 4B, C, movie 2), spleen (Fig. 4G), thymus, liver, skin, heart, and lungs (Fig. S4A). Samples from untreated mice displayed the single unrecombined color code (Fig. S4A, B, S6A). In the small intestine, clonal strings of cells extending from the crypts to the villi tips indicate stable propagation of color codes through cell divisions (Fig. 4B, C, Fig. S4C, D).

**Fig. 4:**
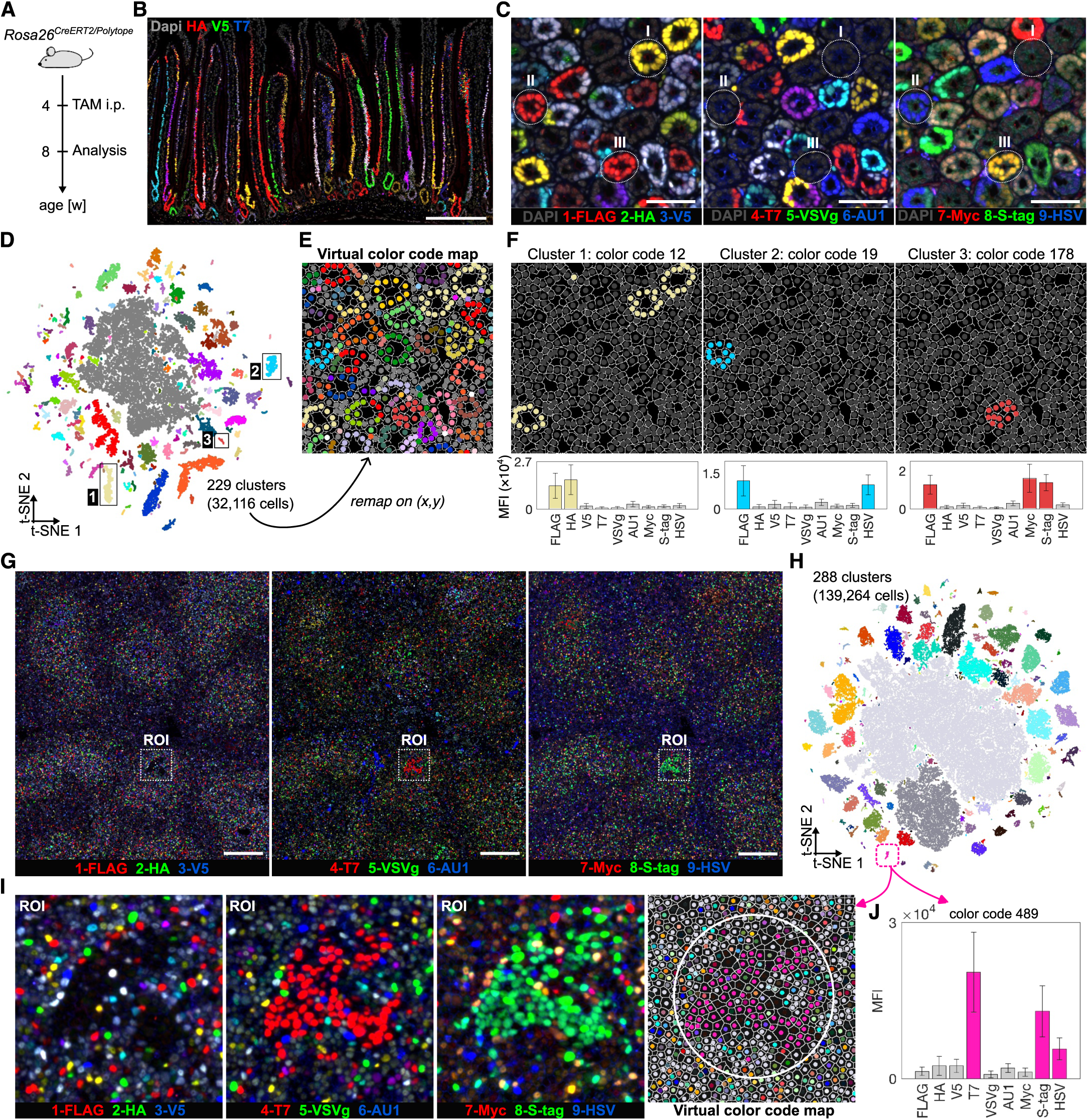
Polytope color coding *in vivo*. (**A**) *Rosa26^CreERT2/Polytope^* mice were treated at four weeks of age with a single dose of tamoxifen via intraperitoneal injection and analyzed after a period of four weeks. (**B, C**) Exemplary multiplexed fluorescent images of villi structures (B, scale bar: 250 µm) or crypts (C, scale bar: 50 µm) in the small intestine. (**D**) t-SNE plot of normalized mean fluorescent intensity values of all nine epitope tag signals derived from multiplexed images of the small intestine. Colors indicate distinct clusters detected by DBSCAN. Boxes specify clusters further analyzed. (**E**) Remapped results of the clustering in (D). Each color indicates a distinct color code. **(F)** Color code maps filtered for specific color code clusters and corresponding mean fluorescent intensity values for each cluster. (**G**) Exemplary multiplexed fluorescent images of the spleen. Scale bar: 200 µm. (**H**) t-SNE plot of normalized mean fluorescent intensity values of all nine epitope tag signals derived from multiplexed images of the spleen, and DBSCAN cluster detection. (**I**) Multiplexed fluorescent images of the region of interest in (G), as well as remapped cluster results from (H). (**J**) Mean fluorescent intensity values of the cluster of interested in (H) and (I).

We next established an image analysis pipeline to extract the color code and xy coordinate for each cell, which is essential for subsequent spatial analyses (Fig. S4E, F). Focusing on a barcoded small intestine sample, we measured the nuclear MFI values of all nine epitope tags for each cell after cellular image segmentation (Fig. S5A). We first attempted to extract color codes by applying thresholding on the bimodal MFI distribution as done before *in vitro*. Comparing this result to a manually annotated dataset (‘ground truth’) showed that the thresholding approach achieves a color code calling accuracy of 80%, likely due to noise and suboptimal signal-to-noise ratio (Fig. S5B, C, E). We hypothesized that a clustering approach might be more robust in detecting cells that share the same color code. Therefore, we aggregated cells of similar MFI values by t-distributed stochastic neighbor embedding (t-SNE) and revealed clusters using Density-Based Spatial Clustering of Applications with Noise (DBSCAN, see Methods for details). Indeed, this approach improved color code calling accuracy to 93%, enabling us to detect 229 distinct clusters across 32,116 intestinal cells (Fig. 4D, S5D, E). Clusters were then mapped onto the xy-plane to create a virtual barcode map (Fig. 4E). Clusters were analyzed for expression of all individual epitopes in order to identify the underlying color code. For instance, cells of cluster 1 harbor the color code ‘12’ (Fig. 4F) and are present in the monoclonal crypt in region I (Fig. 4C). Cells in the distinct clusters 2 and 3 expressed the color codes ‘19’ and ‘178,’ respectively (Fig. 4F), and were located in the monoclonal crypts within regions II and III (Fig. 4C). These results indicate robust color code detection through clustering regarding specificity (distinct color codes visible in the raw data fall into distinct clusters) and sensitivity (same color codes visible in the raw data fall into the same cluster).

We applied this approach to analyze images obtained from the spleen of the same fate-mapping experiment (Fig. 4G, S6B, C, Table S2). We detected 288 distinct color code clusters derived from 139,264 cells (Fig. 4H). As expected, the majority of cells with distinct color codes were distributed randomly throughout the spleen (Fig. 4G). However, we noticed locally confined clones in the white pulp that had emerged in the four weeks chase period. For instance, we identified a dominant CD45^+^B220^+^ B cell clone with the color code ‘489’ in a germinal center (Fig. 4I, J, S6D). In summary, these data demonstrate epitope barcoding *in vivo* as well as reliable identification and mapping of color codes across various tissues, providing insights into clonal dynamics.

## Temporal dynamics of H2B-Polytope turnover after recombination

While the Polytope locus can be recombined at the DNA level and inherited by daughter cells, non-recombined Polytope molecules may persist for some time before they are fully lost. During histone turnover, newly synthesized histones are incorporated into nucleosomes, while ‘old’ histone proteins undergo degradation (*29*). Although this process is typically considered replication-dependent, postmitotic cells, such as cardiomyocytes, also exhibit rapid replication-independent turnover rates (*30*). In the context of Polytope epitope barcoding, ‘old’ H2B-fusion proteins containing the unrecombined Polytope peptide must be replaced by the newly generated H2B-Polytope variant following Cre/loxP recombination (Fig. S7A). Because this time lag is critical for the use of Polytope, we determined the latency from DNA recombination to full replacement of ‘old’ H2B-Polytope proteins with the new variant. To test this, we generated a stable fibroblast cell line derived from *Rosa26^CreERT2/Polytope^* mice and applied a short pulse of 4-hydroxytamoxifen (OHT) *in vitro*, followed by time-course analysis up to seven days post recombination (Fig. S7A). We measured the mean fluorescent intensity of cells after immunostaining with anti-HA-tag antibody (Fig. S7B, C). The data indicate onset of H2B-Polytope protein replacement on day 2 and its completion from day 5 onwards as the percentage of HA-tag-negative cells stabilized at this point (Fig. S7B, C). This kinetic is strictly histone turnover dependent as the DNA recombination pattern was fixed by 24 hours after recombination (Fig. S7D).

To assess the turnover of H2B-Polytope molecules *in vivo*, we administered a single TAM injection to induce recombination in adult *Rosa26^CreERT2/Polytope^* mice and analyzed the tissues after a five-day chase period (Fig. S7E). Consistent with our *in vitro* findings, distinct Polytope color codes were detectable in pancreatic tissue five days post-induction (Fig. S7F). Importantly, in addition to neighboring cells sharing the same color code, we identified individual cells with unique barcodes. The presence of these single cells suggests they did not undergo proliferation during the five-day period, allowing the “old” histone peptide to dilute to background levels independently of cell division. Collectively, these findings highlight a five-day latency period in pulse-chase Polytope barcoding experiments.

## Epitope barcoding reveals local clonal expansion and fragmentation characteristics of fetal liver cells

To further explore the potential of Polytope for deconvolution of the clonal architecture arising in development across embryonic tissues, we ubiquitously labeled cells in developing *Rosa26^CreERT2/Polytope^*mice on embryonic day 8.5 (E8.5). Analysis of whole embryo sections on E16.5 (Fig. S8A) revealed the columnar expansion of neural progenitors in the cerebral cortex and midbrain (Fig. S8B, C), the intermingling of progenitors forming ducts in the kidney, villi structures in the intestine, and branching in the fetal lung (Fig. S8D-F). Additionally, clonal patches were detected in the adrenal gland, pancreas, and olfactory bulb (Fig. S8G-I). These results suggest that Polytope is versatile for conducting developmental studies across diverse organs.

An organ of particular complexity is the developing fetal liver: Hepatocyte development and fetal hematopoiesis progress in parallel between E10.5 and E16.5, making the fetal liver a multilineage and multifunctional organ which eventually changes from hematopoiesis to metabolism around E16.5 (*31*, *32*). We therefore set out to shed light on the clonal composition of the major cell types (hepatocytes and hematopoietic cells), i.e. clone sizes, clone spreading, clone distribution, and peak periods of clonal expansion. We conducted spatial Polytope fate-mapping by unbiased labeling of embryos on E10.5, followed by analysis of livers in E16.5 (Fig. 5A-K)), or adult (Fig. 5L, M) mice. In addition, we induced barcodes on E14.5, followed by adult analysis (Fig. 5L, N). For the fetal liver analysis, we expanded our Polytope antibody panel to include markers for hepatoblasts and hepatocytes (DLK1 and/or CK18), hematopoietic progenitors (Kit), erythroid progenitors (Ter119), macrophages (F4/80), monocytes/granulocytes (CD11b), and other immune cells (CD45), utilizing IBEX multiplexed imaging (Table S2). The liver buds are formed until E10.0 from the foregut by out-migrating hepatoblasts, which start to extensively proliferate (Fig. S8J). At the same time, hematopoietic progenitor cells enter the liver, expand and differentiate, giving rise to nucleated erythroid cells forming erythroblastic islands, macrophages and other immune cells. Across 56,761 fetal liver cells, we identified 108 distinct color code clusters (Fig. 5B), and determined the identity of 93% of these cells by categorizing them into hepatic cells, hematopoietic progenitors, erythroid cells, macrophages, myeloid cells, or other immune cells (Fig. 5C). We determined each cell’s xy-position, cell type, and color code, and projected these data onto a 2D virtual map (Fig. 5D-G). These images suggest cell-type-restricted, microscopic compartmentalization, which might reflect local clonal outgrowth. Neighborhood analysis determines the lineage composition of cells in the vicinity of a reference cell (iterative for all cells) and compares this data to a random distribution. We find that cell types tend to be in close proximity to their kin (Fig. 5H). Prominently, erythroid cells show strong clustering among themselves and are diminished in the neighborhood of other cell types. Hematopoietic progenitors cluster among themselves, but are also associated with hepatic cell neighborhoods. The observed cell type clustering could result from local clonal outgrowth, or, alternatively, from other mechanisms, including selective migration.

**Fig. 5:**
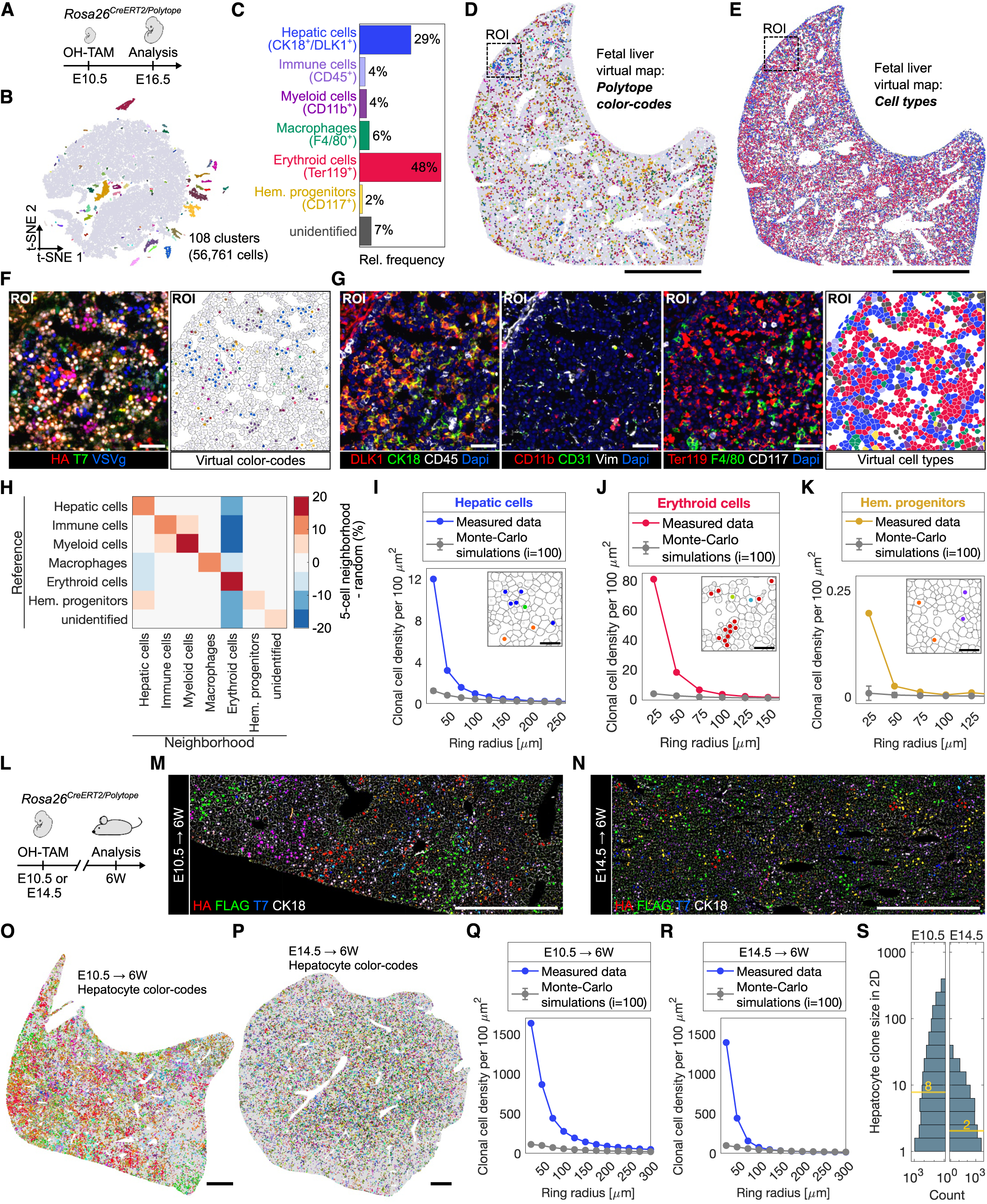
Spatial fate-mapping of the developing fetal liver and from embryonic development to adulthood. (**A-K**) Polytope color coding on the developing fetal liver from E10.5 until E16.5. (**A**) Pregnant mice carrying *Rosa26^CreERT2/Polytope^* embryos were treated with OHT by oral gavage on E10.5 and embryos were analyzed on E16.5. (**B**) t-SNE plot of normalized mean fluorescent intensity values of all nine epitope tag signals derived from multiplexed images of the fetal liver. Colors indicate distinct clusters detected by DBSCAN analysis. (**C**) Quantification of cell types detected, including hepatic cells (DLK1^+^ and/or CK18^+^ hepatoblasts and hepatocytes, blue), erythroid cells (Ter119^+^, red), macrophages (F4/80^+^Ter119^−^, green), hematopoietic progenitors (CD117^+^, yellow), myeloid cells (CD11b^+^F4/80^−^) and remaining immune cells (CD45^+^/CD11b^−^ F4/80^−^). **(D)** Virtual map of color codes and **(E)** cells types (color legend in (C)). Scale bar: 1 mm. (**F**) Exemplary fluorescent image of the region of interest (left) and remapped color code cluster results (right). Scale bar: 50 µm. (**G**) Fluorescent images of cell marker expression and retrieved segmentation and cell type results (far right, color legend in (C)). (**H**) Cell type neighborhood analysis. Frequency of cell types found in the five nearest neighbors normalized to random distribution. Rows indicate reference cell and columns neighboring cells. (**I-K**) Detection of clonal patches and spatial expansion analysis for hepatic cells, erythroid cells and hematopoietic progenitors. Inserted panels depict exemplary clonal patches and spatial distribution. Scale bar: 25 µm. (**L-S**) Polytope color coding of hepatic progenitors from E10.5 or E14.5 until adulthood. (**L**) Experimental design. Recombination in developing *Rosa26^CreERT2/Polytope^* embryos was induced *in utero* at either E10.5 or E14.5 and mice were analyzed at six weeks of age. (**M-N**) Exemplary fluorescent images of the liver from either E10.5 (left) or E14.5 (right) labeled mice. Scale bar: 500 µm. **(O, P)** Remapped color code cluster results and (**Q, R**) spatial clonal distribution analysis. Scale bar: 1 mm. (**S**) Quantification of 2D hepatocyte clone sizes based on color code and spatial separation.

To resolve this question, we analyzed Polytope color codes using a ring analysis approach (*33*). This method quantifies the density of same color-coded cells in concentric rings of increasing diameter and compares the data to one hundred iterations of Monte Carlo simulations to determine whether observed clustering is actual or random. This approach further provides information about the average spatial expansion of clones, marked by the distance at which the measured data converges with the Monte-Carlo simulations. Applying this analysis to hepatic cells demonstrates significant clustering of clones compared to random distribution, with clonal expansion spanning up to 200 µm in diameter (Fig. 5I). Closer visual inspection shows that while hepatic cells of the same clone can be direct neighbors in close proximity, we also observed clone fragmentation over a diameter of up to several hundred micrometers (Fig. S9A, B). Hepatic patches (clusters of hepatocytes) are not monoclonal, and rather, intermingling of different hepatic clones is observed (Fig. S9B). Erythroid cells display clonal clustering and fragmentation, although with stronger clustering behavior (Fig. 5J, S9C, D), while hematopoietic progenitors exhibit a more scattered clonal organization (Fig. 5K, S9E, F).

Together, these findings indicate that fetal liver growth between E10.5 and E16.5 is driven by the local expansion of intermingling hepatic clones with clone sizes of up to 20 cells measured in 2D (Fig. S9G). Despite being concentrated in one location, clones are fragmented. Erythromyeloid progenitors that enter the liver at E10.5 give rise to large erythroid clones of up to 60 cells in 2D (Fig. S9H), while hematopoietic progenitors form smaller local clones of up to five cells in 2D. Overall, resolving clones by Polytope barcoding demonstrated that local clonal expansion and clone fragmentation are hallmarks of fetal liver development between E10.5 and E16.5.

## Timed barcoding reveals key phases of clonal expansion of fetal liver cells

The liver bud undergoes a period of accelerated growth between E10.5 and E15.5 from approximately 0.4 mg to 41.6 mg in weight (∼100-fold, Fig. S10A). This increase in weight continues from E15.5 until adulthood (∼42-fold). As hepatoblasts expand in numbers, they simultaneously begin to undergo lineage specification, acquire location-specific functions and develop polyploidy, while increasing cellular volume (*31*). During this period, hematopoietic cells are progressively lost. This raises the question of the main drivers of liver growth throughout development, which may include clonal expansion or other mechanisms, such as increases in cellular volume. To address this question, we applied timed Polytope barcoding and clonal analysis by labeling fetal liver cells at either E10.5 or E14.5 in developing *Rosa26^CreERT2/Polytope^* embryos followed by adult analysis using IBEX multiplexed imaging (Fig. 5L-N, S10B, Table S2). Focusing on hepatocytes only, we retrieved in total 161 or 146 distinct color codes, respectively (Fig. S11A, S12A, Fig. 5O, P). Ring analysis demonstrated local expansion of hepatocyte clones in E10.5-treated livers, with clones reaching diameters of up to 300 to 600 µm. In contrast, hepatocyte clones in E14.5-treated livers were smaller, typically under 150 µm in diameter, indicating locally confined outgrowth (Fig. 5Q, R). We analyzed color codes further, by focusing on their spatial distribution. Using spatial clustering to separate independent clones sharing the same color code (Fig. S11B-H, S12B-H), we compared clone sizes on 2D liver sections, resulting in an incomplete, but comparable 2D measurement. This analysis showed a 0.26-fold decrease in mean clone size between E10.5 and E14.5 (Fig. 5S). Clones in E10.5 barcoded livers comprised up to several hundred nuclei in 2D, while those in E14.5 barcoded livers were smaller, typically fewer than 40 nuclei (Fig. 5S). These findings suggest that liver development occurs in two distinct phases. First, massive clonal expansion of hepatoblasts between E10.5 and E14.5 marks a proliferative stage driving liver bud growth. Second, following the onset of metabolic activity and the formation of hepatic sinusoids after E14.5, cells become less proliferative, indicating that clonal expansion is no longer the dominant driver of liver growth.

## Discussion

The Polytope system presented in this study marks an advancement in fate-mapping technologies by integrating the spatial resolution of imaging-based methods with the labeling complexity of barcoding technologies. Our *in vitro* and *in vivo* experiments demonstrate robust functionality of Polytope for clonal analysis across various tissues. Spatial fate-mapping results of liver development illustrate the system’s ability to reveal key aspects of tissue organization, including local expansion and fragmentation of clones. Timed barcoding experiments further uncovered the temporal changes of clonal expansion during liver development.

With an increased label diversity of up to 512 distinct color codes generated through random recombination of epitope tags, Polytope allows for precise tracing of individual clones *in situ*. Achieving such high label diversity enables the resolution of stem cell systems in densely packed progenitor regions, such as the small intestine, where approximately 14±2 Lgr5^+^ stem cells are found per crypt (*9*). Polytope further improves tracing of clones over several hundred micrometers, accounting for situations in which clonal cells separate due to migration or clonal fragmentation – a phenomenon observed in embryonic development as well as hematopoiesis (*34*) and neurogenesis (*35*), where daughter cells actively migrate from the stem cell niche to distant regions. In a recent study, we applied Polytope to map the developmental pathways of macrophage progenitors giving rise to Kupffer cells, revealing local seeding and expansion dynamics as well as quantifying clone sizes (*36, in preparation*).

When combined with extended antibody panels in multiplexed imaging experiments, Polytope not only captures each cell’s clonal history and spatial coordinates but also provides insights into cell type identity (e.g., CD8) and cellular states (e.g., PD-1). This multidimensional data offers a comprehensive view of tissue organization and spatio-temporal clonal dynamics, and enables direct observation of differentiation pathways within the tissue. As Polytope color codes are obtained via multiplexed imaging, readout costs (approx. 50€ per slide when using IBEX multiplexed imaging) are reduced compared to other spatial barcoding technologies relying on sequencing. The methods presented here make Polytope widely accessible, requiring only a standard wide-field fluorescent microscope for color code readout without the need for specialized equipment. Additionally, as the barcode information is present on the DNA, RNA and protein level, Polytope has the potential to be integrated with spatial transcriptomics and proteomics in the future, opening opportunities to examine spatial OMICs datasets in parallel with lineage tracing. In summary, spatial fate-mapping with Polytope provides deep insights into clonal architectures, making it a valuable tool for exploring fundamental principles of cellular systems within their native microenvironments (Fig. S13).

In summary, Polytope offers detailed insights into stem cell systems by deconvolving cellular lineages at both clonal and spatial resolution within their native microenvironment (Fig. S13).

## Materials and Methods

### Molecular cloning

The Polytope cassette was assembled in two steps by In-Fusion cloning (Takara Bio) of synthesized double-stranded DNA fragments (Eurofins Genomics) and cloned into a SalI/EcoRV linearized AAVS1-CAG vector (AAVS1-CAG-hrGFP was a gift from Su-Chun Zhang; Addgene plasmid #52344), generating the AAVS1-CAG-H2B-Polytope plasmid. The complete sequence is available in the supplementary materials (Table 3). The Polytope cassette was further cloned into a Rosa26 vector backbone. In order to target the AAVS1 locus, the gRNA sequence 5’-GGGGCCACTAGGGACAGGAT-3’ was cloned into the pSpCas9(BB)-2A-Puro (PX459) V2.0 plasmid (this plasmid was a gift from Feng Zhang; Addgene plasmid #62988), generating the pSpCas9-gRNA(AAVS1) plasmid.

### CRISPR/Cas9 knock-in into AAVS1 locus in HEK cells

HEK293 cells were maintained in Dulbecco’s modified Eagle’s medium (DMEM) (Thermo Fisher Scientific) supplemented with 10% heat-inactivated fetal calf serum (FCS) (Thermo Fisher Scientific) and 1% penicillin-streptomycin–glutamine solution (Thermo Fisher Scientific). Transfection with the donor construct (AAVS1-CAG-H2B-Polytope) and the pSpCas9-gRNA(AAVS1) plasmid was conducted with Lipofectamine 2000 (Invitrogen) according to supplier’s protocol. Knock-in cells were selected with 2.5 µg/mL puromycin starting 2 days after transfection for 7 days, before cells were seeded at low density and clones were handpicked from the dish. Established knock-in cell lines were genotyped using DPPrim_073: 5’-CCAGCTCCCATAGCTCAGTC-3’, DPPrim_075: 5’-ACAGGAGGTGGGGGTTAGAC-3’ and DPPrim_111: 5’-TGCCAAGGCAAAGGAGACC-3’ and a heterozygous knock-in cell line was selected for further experiments.

### Cellular *in vitro* assay of Polytope color coding and IBEX multiplexed imaging

*AAVS1^Polytope/+^* cells were treated for 16h with 100, 200 or 300 U/mL tat-Cre (Merck) in complete medium and propagated for five days, before being seeded on IBIDI 8-well imaging slides coated with poly-D-lysine (Gibco). Two days later, cells were washed with PBS, fixed for 15 minutes in 4% PFA/PBS, followed by washing in PBS, solubilization in PBS containing 0.1% Triton X-100 (PBST) for 10 minutes at room temperature and blocking in PBST + 1% BSA + 10% donkey serum for one hour at room temperature. A complete list of the antibody panel for each experiment is provided in Table 2 and multiplexed imaging was performed as described in the manual protocol of IBEX (*26*). Briefly, directly conjugated antibodies (Alexa Fluor 488 or 555 or 647), were incubated at 4°C over night in staining buffer (PBST + 1% BSA) with subsequent washing with staining buffer, PBST, incubation with 1 µg/mL Dapi/PBST and washing with PBS, before mounting in a mixture of Fluoromount-G/PBS (50%/50%). Pictures of 8-well imaging slides were captured on a bright-field fluorescent microscope (Keyence, BZ-X810) using a 20X Plan Apochromat objective (NA: 0.75, WD: 0.6 mm), followed by image stitching with the included software. Wells were washed three times with PBS and samples were bleached in 1.0 mg/mL LiBH_4_ solution (dissolved in diH2O) for 20 min at room temperature under exposure to a white LED light (commercial daylight lamp). Samples were washed three times in PBS and incubated in blocking buffer for 30 min at room temperature before the next IBEX cycle started. Effective bleaching after each cycle was confirmed by fluorescent microscopy.

### Generation of *Rosa26^Polytope/+^* mice

The *Rosa26^Polytope/+^* mouse model (C57BL/6JRj-Gt(ROSA)26Sor^tm7Nki^ MGI:7339240; synonym: C57BL/6JRj-Gt(ROSA)26Sor^tm7(CAG-polytope3.0)Nki^) was generated by homology directed recombination in embryonic stem cells derived from C57BL/6JRj mice, which were further injected into C57BL/6 blastocysts at E3.5 days post conception (dpc). Chimeric males were backcrossed to C57BL/6J females to transmit the *Rosa26^Polytope/+^* knock-in allele through the germline. Obtained mice were further analyzed by Southern blotting using a biotinylated 820-bp probe located upstream of the first exon and the short arm of homology (*37*). The probe was generated by PCR amplification from genomic C57BL/6 DNA using the primers #2,424: 5′-GCAAAGGCGCCCGATAGAATAA-3′ and #2,425: 5′-CCGGGGGAAAGAAGGGTCAC-3′. Probe labelling, hybridization and detection were done with the North2South Biotin Random Prime Labelling Kit, and the North2South Chemiluminescent Hybridization and Detection Kit (Thermo Scientific). Heterozygous *Rosa26^Polytope/+^*as well as homozygous *Rosa26^Polytope/Polytope^* mice are viable and fertile, and do not show any obvious phenotype. Mice were genotyped using primers DPPrim_299: 5’-AAGGGAGCTGCAGTGGAGTA-3’, DPPrim_300: 5’-TAAGCCTGCCCAGAAGACTCC-3’, DPPrim_301: 5’-GGCGTTACTATGGGAACATACG-3’. Mice were kept in individually ventilated cages under specific pathogen-free conditions in the animal facility at the German Cancer Research Center (DKFZ, Heidelberg) and Netherlands Cancer Institute (NKI, Amsterdam). Male and female mice were used, no randomization was done, no blinding was done and no animals were excluded from the analysis. No statistical methods were used to predetermine sample size. All animal experiments were performed in accordance with institutional and governmental regulations, and were approved by the Regierungspräsidium (Karlsruhe, Germany) or the Animal Welfare Committees of the Netherlands Cancer Institute.

### Histopathology

Systematic histopathological evaluation was performed on mice at 9 and 15 weeks of age. Littermates of wildtype, heterozygous and homozygous genotype were generated through breedings with heterozygous parents. Per genotype, 6 littermates comprising 3 females and 3 males were used for analysis. A pathologist evaluated internal organs by macroscopic evaluation at necropsy, and by microscopic evaluation of hematoxylin-eosin-stained internal organs.

### Fluorescent-activated cell sorting (FACS), Polytope PCR, Sanger sequencing and sequence alignment of Polytope HEK cells

Tat-Cre treated HEK *AAVS1^Polytope/+^* cells were harvested, transferred into FACS buffer (PBS + 5% FCS), containing Sytox blue and live, single cells were sorted into wells of a 96-well plate containing complete medium as described above. Cells were allowed to expand for 14 days with media exchange every 3 days, and successfully outgrown clones were analyzed by full-length Polytope PCR (DPPrim_281: 5’-GTCCGAGGGTACTAAGGCCATC-3’, DPPrim_282: 5’-TGGTATTTGTGAGCCAGGGCATT-3’) and subsequent Sanger sequencing using the same primers, as well as IBEX multiplexed imaging as described above. Sequence alignment of epitope tag cassettes on the obtained Sanger sequencing .fas files was performed using the Snapgene software.

### *In vivo* induction of Polytope recombination

Homozygous *Rosa26^Polytope/Polytope^* mice (C57BL/6JRj-Gt(ROSA)26Sor^tm7Nki^ MGI:7339240) were crossed to homozygous *Rosa26^CreERT2/CreERT2^* (B6.129-Gt(ROSA)26Sor^tm1(cre/ESR1)Tyj/J^) mice to generate *Rosa26^Polytope/CreERT2^* animals. Mice at four weeks of age were injected intraperitoneally with 80 µg/g body weight tamoxifen solved in peanut oil. The tamoxifen stock solution was prepared by dissolving 1 g tamoxifen (Sigma) in 4 ml absolute ethanol and 36 ml peanut oil (Sigma) at 55 °C.

### *In utero* induction of Polytope recombination

Timed matings were set up between *Rosa26^Polytope/Polytope^* and *Rosa26^CreERT2/CreERT2^* mice. Eight, ten or fourteen days after the day of the plug (day 0.5), pregnant mice were treated by oral gavage with a single dose of 50-60 µg/g body weight 4-hydroxytamoxifen (Sigma) to induce barcoding in the developing embryos at either E8.5, E10.5 or E14.5, together with 1.25 mg progesterone (Sigma) to sustain pregnancy. The pups were delivered on E20.5 by caesarean section, raised by foster mothers and analyzed at 6 weeks of age; or dissected from the pregnant mother at E16.5 and analyzed at the fetal stage.

### Sample collection, freezing, cryosectioning and IBEX multiplexed imaging

For collection of organs from adult mice, euthanized mice were perfused via the left ventricle at low flow rate with 10 mL of Hank Balanced Salt Solution (HBSS) containing 20 U/mL heparin (Sigma) with the right atrium opened through incision for outflow. Organs were collected and placed in cold PBS on ice, before being fresh frozen in optimal cutting temperature (O.C.T) compound (Tissue-Tek) using liquid nitrogen-chilled 2-methylbutane. For collection of embryos, fetuses were directly dissected from euthanized mice and placed in cold PBS before being frozen in O.C.T. compound as described above. Cryoblocks were stored at -80°C. Blocks were cut as 5 µm sections on an Epredia CryoStar NX50 cryostat and placed on SuperFrost Ultra Plus GOLD glass slides (Epredia), before being stored at -80°C after a 1h drying period at room temperature. Before immuno-staining, sections were dried for 30 min at room temperature, excess O.C.T. was removed, the sample was encircled with a barrier pen, and 4% PFA/PBS was added for 15 minutes at room temperature, before three consecutive washing steps in PBS, and subsequent solubilization in PBST (PBS + 0.1% Triton X-100) for 10 min at room temperature and blocking in blocking buffer (PBST + 1% BSA + 10% donkey serum) for 1h at room temperature. Due to high autofluorescence, embryo sections were pre-bleached using the TrueVIEW autofluorescence quenching kit (Vector Laboratories) according to supplier’s protocol after fixation, but before solubilization and blocking. A complete list of the antibody panel is provided in supplementary table 2 and multiplexed imaging was performed as described in the manual protocol of IBEX (*26*). Briefly, directly conjugated antibodies, were incubated at 4°C over night in staining buffer (PBST + 1% BSA) with subsequent washing with staining buffer, PBST, incubation with 1 µg/mL Dapi/PBST and washing with PBS, before mounting in Fluoromount-G/PBS (50%/50%). Whole section images were either acquired on a Keyence BZ-X810 (20X objective/ 0.75 plan-apochromat) or a Zeiss Axioscan 7 (20X objective/ 0.8 plan-apochromat air). Coverslips were removed by demounting in PBS, sections were washed in PBS and photobleached in 1.0 mg/mL LiBH_4_ solution (dissolved in diH2O) for 20 min at room temperature under exposure to a white LED light (commercial daylight lamp). Samples were washed three times in PBS and incubated in blocking buffer for 30 min at room temperature before the next IBEX cycle started. Effective bleaching was confirmed on a Keyence BZ-X810 (20X objective/ 0.75 plan-apochromat) widefield fluorescent microscope.

### Culture and treatment of fibroblasts derived from *Rosa26^CreERT2/Polytope^* mice

Ears and tails from *Rosa26^CreERT2/Polytope^* mice were sterilized in 70% ethanol for 30 seconds, washed in sterile PBS, cut into small pieces and placed into 15 mL tubes containing 10 mL sterile DMEM (without serum) supplemented with 0.5 mg/mL Liberase DL (Roche). Digestion was performed at 37°C for 90 minutes while shaking, followed by centrifugation at 300g, resuspension of the tissue pieces in 10 mL DMEM supplemented with 10% FSC and 1% penicillin-streptomycin–glutamine solution, and transfer to a 10 cm CELLSTAR cell culture dish (Greiner). Tissue pieces were cultured for five consecutive days, allowing the fibroblasts to migrate out of the tissue pieces and adhere to the culture dish. Cells and tissue pieces were collected after trypzination and fibroblasts were isolation by passing the suspension through a 40 µm strainer and further propagated. Fibroblasts were immortalized via transduction of the SV40 LT gene (pBABE-puro SV40 LT was a gift from Thomas Roberts; Addgene plasmid #13970) using ecotropic gamma-retrovirus produced in Plat-E cells as described before (*38*). Cells were selected with 2.5 µg/mL puromycin three days after transduction and further propagated. Recombination was induced by a short pulse of 1h with 400 mM 4-hydroxy-tamoxifen (Sigma) added to the media. Treated *Rosa26^CreERT2/Polytope^* fibroblasts as well as untreated controls were analyzed 1-7 days after pulse by harvest, washing in PBS, fixation in 4% PFA for 15 minutes, washing in PBS, and storage in PBS containing 0.1% Triton X-100 at 4°C until further analysis. Once all samples were collected, fixed cells were incubated in blocking buffer (PBS containing 0.1% Triton X-100 and 1% BSA) for 1h at room temperature, before immunostaining with 1:600 HA-Tag (C29F4) Rabbit mAb Alexa Fluor-555 (CST, #55420) over night at 4°C. Stained samples were washed twice with FACS buffer (PBS + 5% FCS), before being analyzed on a FACSAriaIII cell sorter (BD Biosciences) in order to measure fluorescent intensities as well as sort 50,000 cells per sample into a 1.5 mL tube containing FACS buffer. Collected samples were washed in PBS and lysed in 25 µl lysis buffer (containing 12.6 µg proteinase K (Thermo Fisher Scientific) in PCR buffer 1 (Roche)). Lysis was performed by incubating cells for 1 h at 55 °C, followed by 95 °C for 10 min to terminate lysis, and cooling to 4 °C before adding the remaining PCR reagents to a final volume of 50 µL. The whole Polytope cassette was amplified by PCR (Expand Long Template PCR System (Roche, cat. no. 11759060001)) using DPPrim_281: 5’-GTCCGAGGGTACTAAGGCCATC-3’, DPPrim_282: 5’-TGGTATTTGTGAGCCAGGGCATT-3’. PCR products were analyzed by gel electrophoresis.

### Computational analysis of microscopy images, color code retrieval and data analysis

Images derived from IBEX multiplexed imaging were registered based on the Dapi channel of each immunostaining cycle (with the Dapi channel of the first imaging round as reference) using the MultiStackReg plugin in Fiji/ImageJ (*Busse & Miura* (*2016): <u>MultiStackReg</u>. Github*, (*39*). Background subtraction was performed using median filtering or by subtracting a beforehand obtained background image (prior to immunostaining).

For data obtained from *in vitro* experiments, nuclear segmentation was performed by global thresholding (Li algorithm) followed by Watershed separation, before measuring mean fluorescent intensity values for each cell, which were further analyzed in MATLAB. The threshold for gating into either positive or negative for each individual tag was obtained by automatically determining the global minimum between the two maxima of the bimodal MFI distribution.

For data obtained from postnatal *in vivo* barcoding experiments, segmentation was performed using pretrained models of Cellpose (intestine: tissuenet 3, spleen: tissuenet 2), using both nuclear (Dapi) and membrane marker (intestine: Na/K-ATPase and CD45, spleen: CD45), followed by mean fluorescent intensity measurement and subsequent analysis in MATLAB (*40*). Color code calling was performed by t-distributed stochastic neighbor embedding (euclidean distance, Barnes–Hut approximation, exaggeration = 4, perplexity = 30, learn rate = 500) on normalized mean fluorescent intensity values, reducing the multidimensional dataset to 2D, followed by color code cluster detection by applying the density-based clustering non-parametric algorithm (DBSCAN, minimum number of neighbors for core point = 5, optimal epsilon value was determined by applying the ‘elbow’ method on k-distance graph, (*41*)).

For testing color code calling accuracy, a dataset of 508 intestinal cells was manually annotated, generating a ‘ground truth’ color code dataset, which was compared to the global thresholding and clustering approach.

For data obtained from *in utero* barcoding experiments followed by embryonic analysis, segmentation was performed by applying the pretrained Cellpose tissuenet 3 model (*40*) based on nuclear (Dapi) and ubiquitous membrane marker (Na/K-ATPase). Nuclear Polytope epitope tag signals were measured for each cell. Cell marker signals were binarized using automated global thresholding (MATLAB), and pixel values were measured by applying the segmentation mask derived from Cellpose. Cell type calling was achieved on binary channel readout. Note that central macrophages in erythroblastic island are not detected as macrophages due to overlapping signals, hence classification as macrophage in this dataset is limited to F4/80^+^ cells outside of Ter119^+^ erythroblastic islands. Color code calling was determined as described above (t-SNE: euclidean distance, Barnes–Hut approximation, exaggeration = 4, perplexity = 30, learn rate = 700; DBSCAN: minimal points = 5, epsilon (determined by elbow method) = 0.95). Neighborhood analysis was performed by determining frequency of cellular identity of the five closest neighbors compared to random distribution. Ring analysis was performed as described previously (*33*) with adjustments to radii as indicated in the figures. Planar clone sizes were determined by applying DBSCAN on xy coordinates for each detected color code (minimal points = 1, epsilon = 100 µm). For data obtained from *in utero* barcoding experiments followed by adult analysis, a segmentation mask was created based on the DAPI channel by global thresholding (Li) and Watershed separation. Cell marker signals were binarized using automated global thresholding (MATLAB). Color code as well as cell type calling was achieved as described above. 2D clone sizes were determined by applying DBSCAN on xy coordinates for each detected color code (minimal points = 1, epsilon = 200 µm).

## Acknowledgments

We thank all members of the van Rheenen and Rodewald laboratory for ongoing support and discussions; Amber Schonk and Svenja Kösegi for experimentation during early development of the Polytope system; Lennart Kester for scientific discussions during conceptualization; Martijn van Baalen for expert flow cytometry; Jeroen Lohuis for mouse handling and support during the characterization and protocol establishment of the Polytope mouse; Günter Küblbeck for technical assistance and expertise; the bioimaging core facility of NKI and the light microscopy core facility of DKFZ; Juliana Kenski and Robin Thiele for scientific discussions; the teams of animal caretakers at NKI and DKFZ for support and expert animal husbandry.

## Funding

D.P. was supported by the Helmholtz Graduate School for Cancer Research fellowship. L.F. was supported by the I&I Helmholtz Initiative (ZT-0027) and the Christiane Nüsslein-Volhard Foundation fellowship. H.-R.R. was supported by the European Research Council Advanced Grant 742883 and the Leibniz program of the Deutsche Forschungsgemeinschaft. J.v.R. was supported by the Ammodo Science Award, the Doctor Josef Steiner Foundation, the Netherlands Organization for Scientific Research (NWO, Gravitation programme IMAGINE!, project number 24.005.009 and VICI project number 09150182110004).

## Author contributions

D.P. conceived the study, designed and performed experiments, interpreted results, and wrote the paper. C.E.J.P. and P.K. generated the *Rosa26^Polytope^* mouse line. L.F. provided support during animal experiments and C-section, and edited the paper. T.v.L. performed the five-day chase experiment. H.A.M. implemented histopathological analysis and edited the paper. J.v.R. and H.-

R.R. supervised the study and edited the paper. The manuscript was approved by all authors.

## Competing interests

The authors declare no competing interests.

## Data and materials availability

The data sets generated and/or analyzed during the current study are available from the corresponding authors on reasonable request.

---

**Suppl. Fig. 1:**
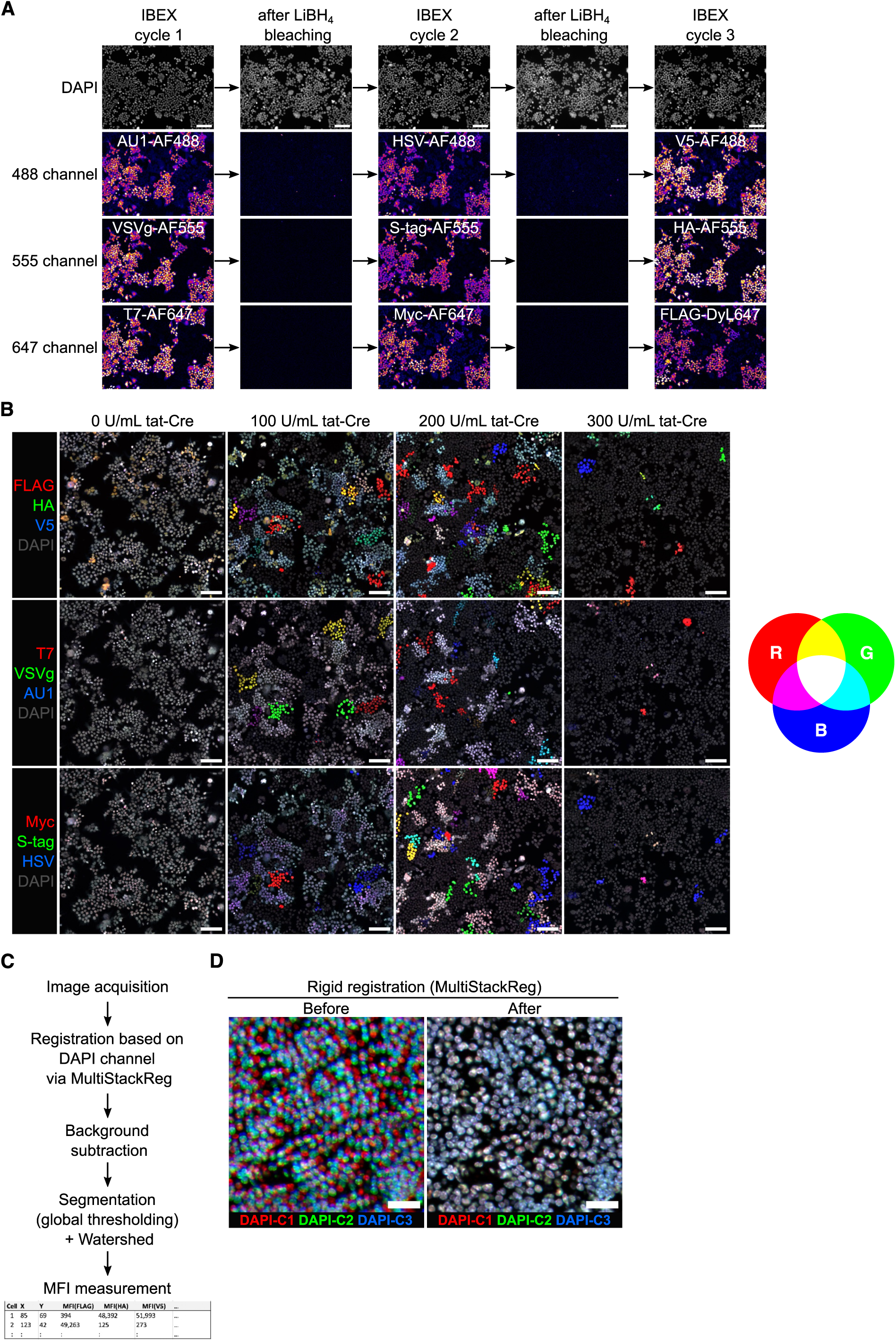
Polytope color code induction *in vitro*, IBEX multiplexed. (**A**) Multiplexed IBEX imaging with all nine Polytope epitope tags, including intermediate images after bleaching. (**B**) Representative images of tat-Cre-treated Polytope cells at varying recombination rates (RGB color system). Scale bar: 100 µm. **(C)** Image processing and analysis pipeline. (**D**) Registration result applying MultiStackReg (see Methods).

**Suppl. Fig. 2:**
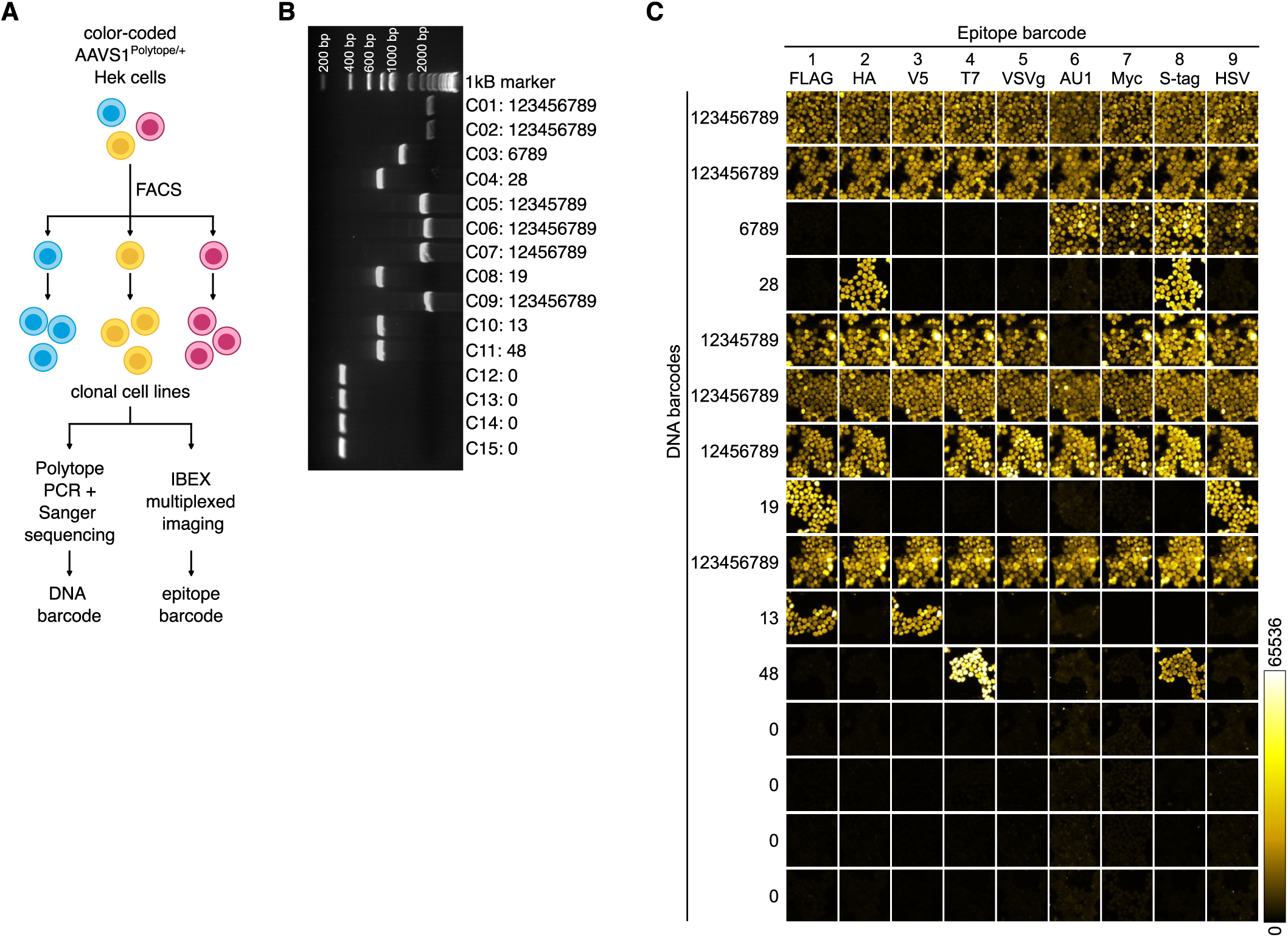
Comparison of the color code detected by multiplexed imaging versus the underlying DNA barcode detected by Sanger sequencing. (**A**) Schematic depicting the generation of clonal cell lines from tat-Cre treated *AAVS1^Polytope/+^*HEK293 cells and the subsequent analysis by Polytope-specific PCR and Sanger sequencing, and IBEX multiplexed imaging. (**B**) Gel electrophoresis of Polytope PCR fragments derived from clonal cell lines. PCR products were further analyzed by Sanger sequencing. **(C)** DNA barcode vs. epitope color code: The numbers on the left represent the DNA barcode derived from sequence alignment, while columns show images from the multiplexed imaging. Each row represents an individual clonal cell line.

**Suppl. Fig. 3:**
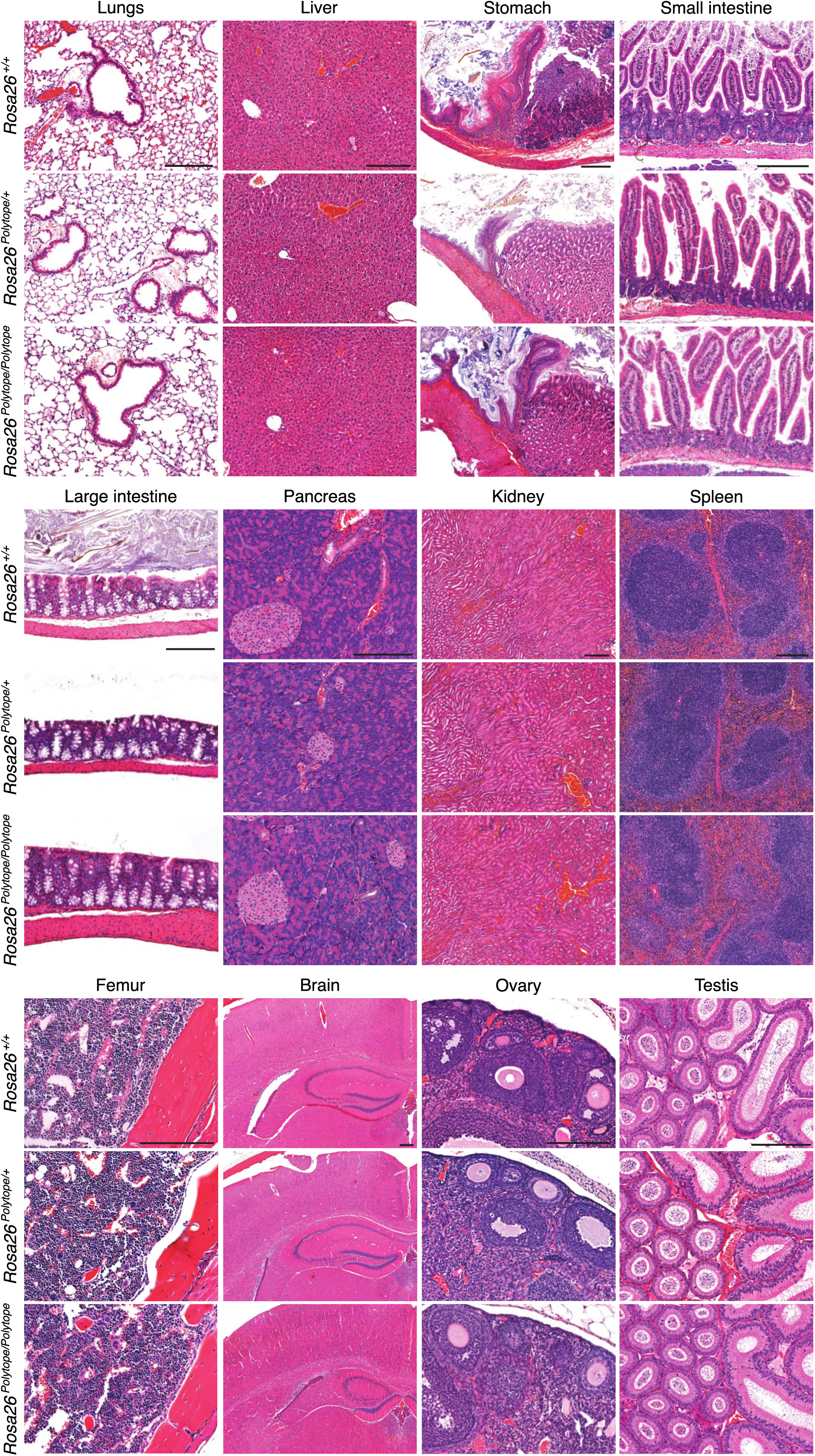
Histopathology of wildtype, heterozygous and homozygous mice. Hematoxylin and eosin staining of the indicated organs. Shown are representative images of 6 mice per genotype, 3 females and 3 males each. All scale bars 200 µm.

**Suppl. Fig. 4:**
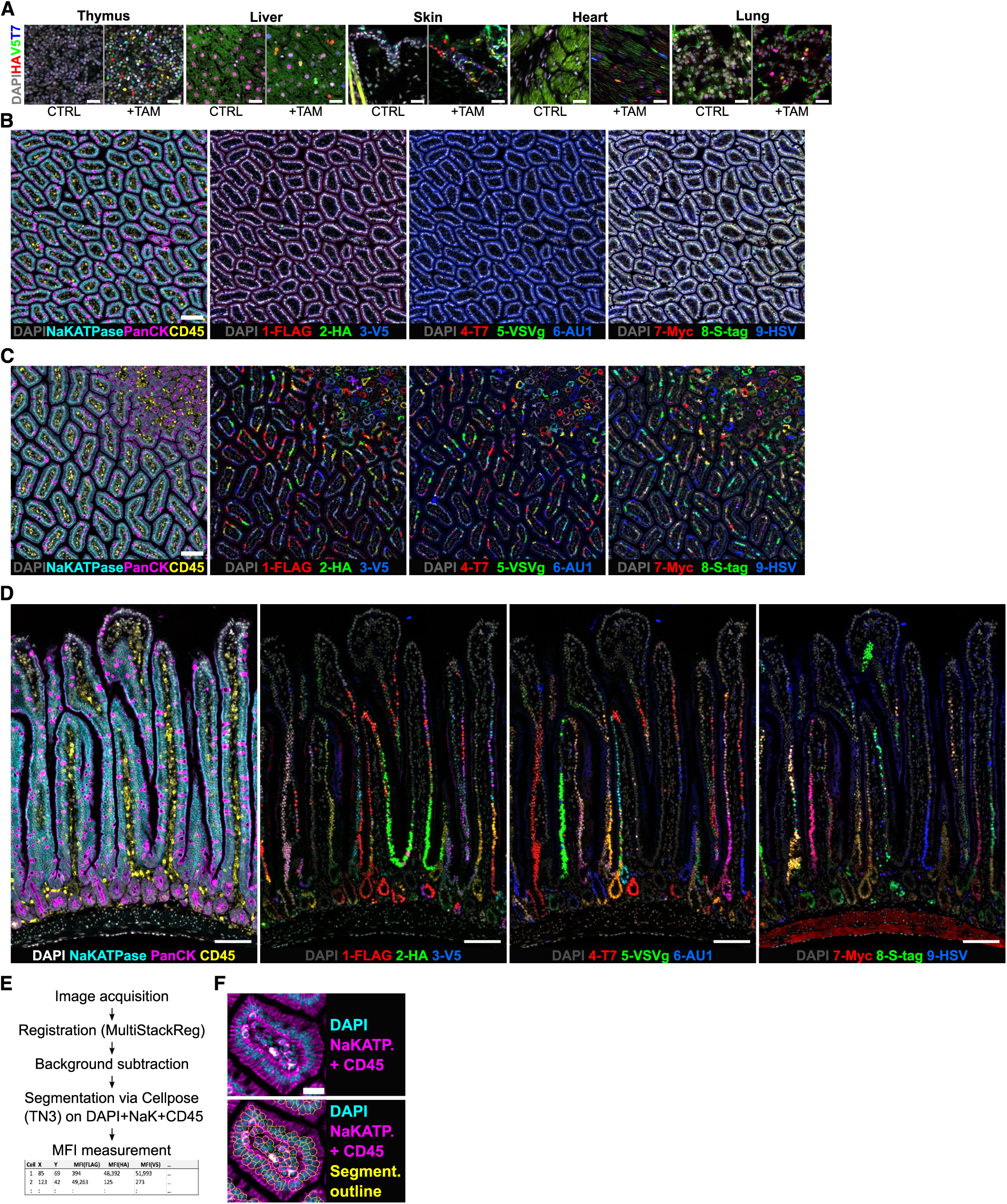
Polytope color coding *in vivo* and image analysis of the small intestine. (**A**) Color code induction in various tissues. (**B**) Multiplexed fluorescent images of the untreated or (**C, D**) TAM-treated small intestine (RGB color system). (**E**) Image analysis pipeline. (**F**) Cellpose TN3 segmentation results (yellow outlines) based on nuclear (DAPI) and membrane (Na/K-ATPase and CD45) marker.

**Suppl. Fig. 5:**
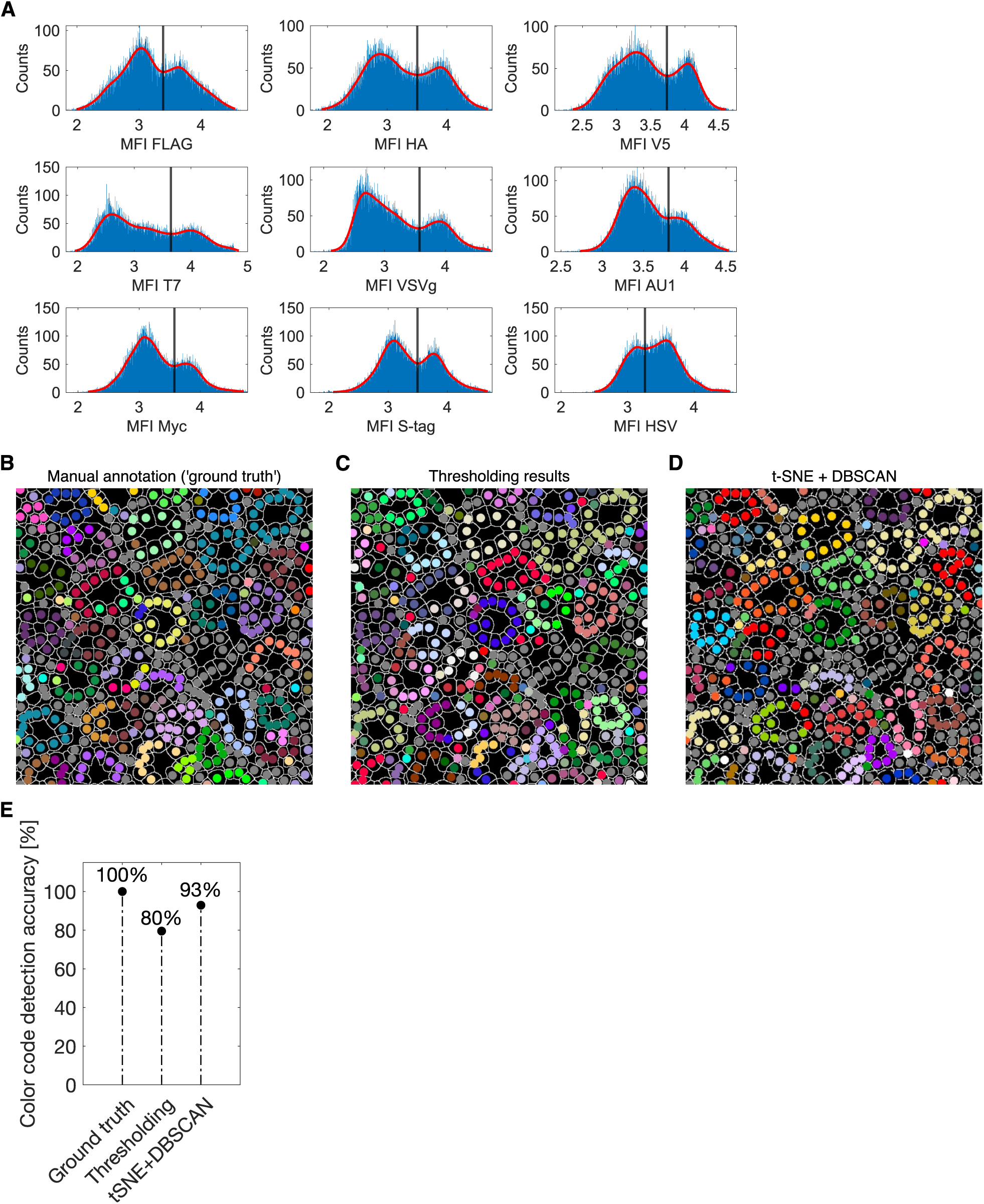
Color code calling of small intestine data. (**A**) Mean fluorescent intensity value histograms of barcoded small intestine samples. The vertical line depicts the threshold determined by automatic segregation of the bimodal distribution. (**B-D**) Virtual color code maps of a manually annotated dataset (‘ground truth’, B), resulting from global thresholding (C) or derived from clustering (t-SNE+DBSCAN, D). (E) Color code detection accuracy of different calling approaches.

**Suppl. Fig. 6:**
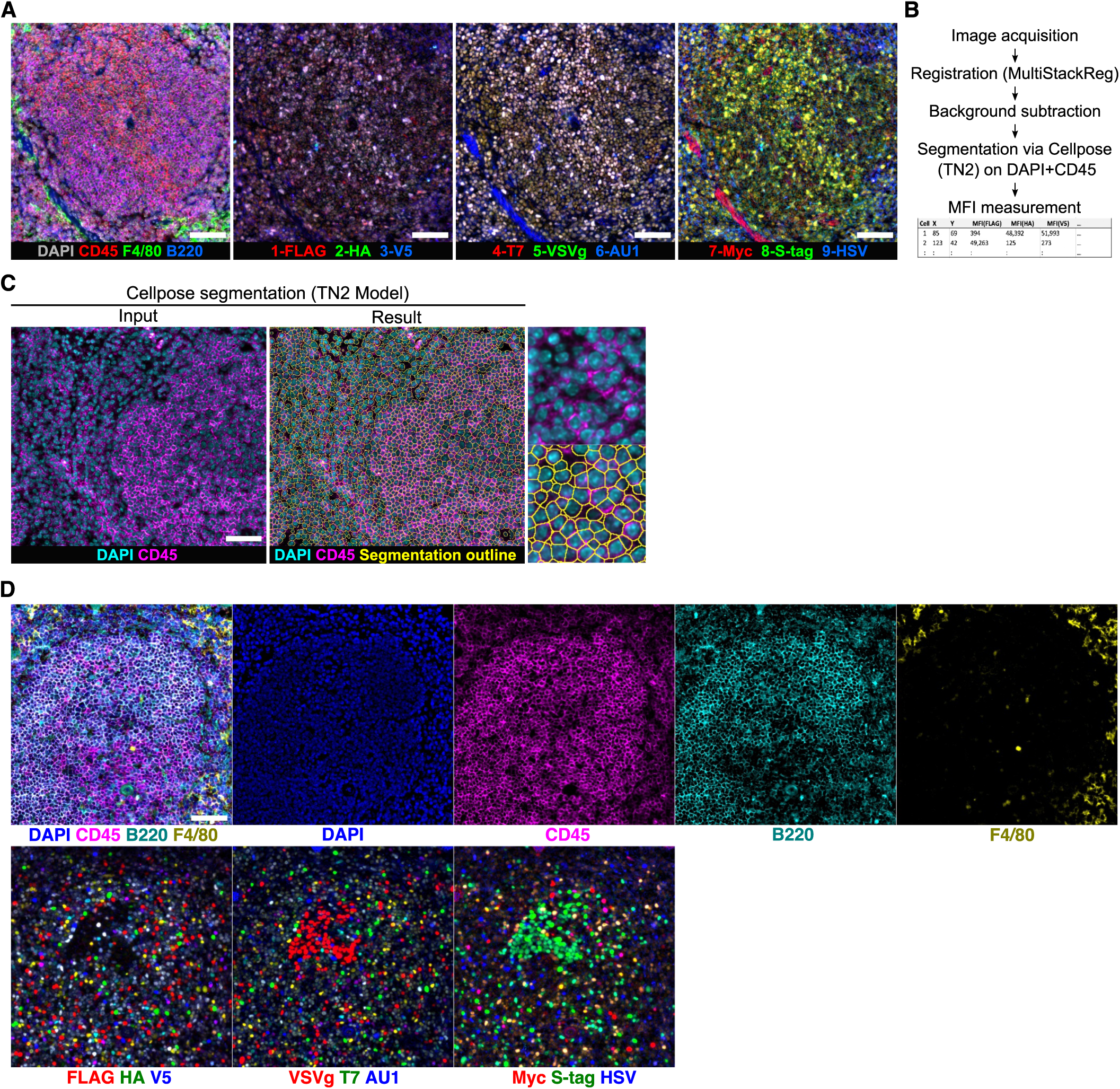
Polytope color coding *in vivo* and image analysis of the spleen. (**A**) Polytope multiplexed imaging of the spleen from an untreated mouse. (**B**) Image analysis pipeline. (**C**) Cellpose TN2 segmentation results (yellow outlines) based on nuclear (DAPI) and membrane (CD45) marker. (**D**) IBEX multiplexed images using the full panel to detect germinal centers as well as Polytope color codes.

**Suppl. Fig. 7:**
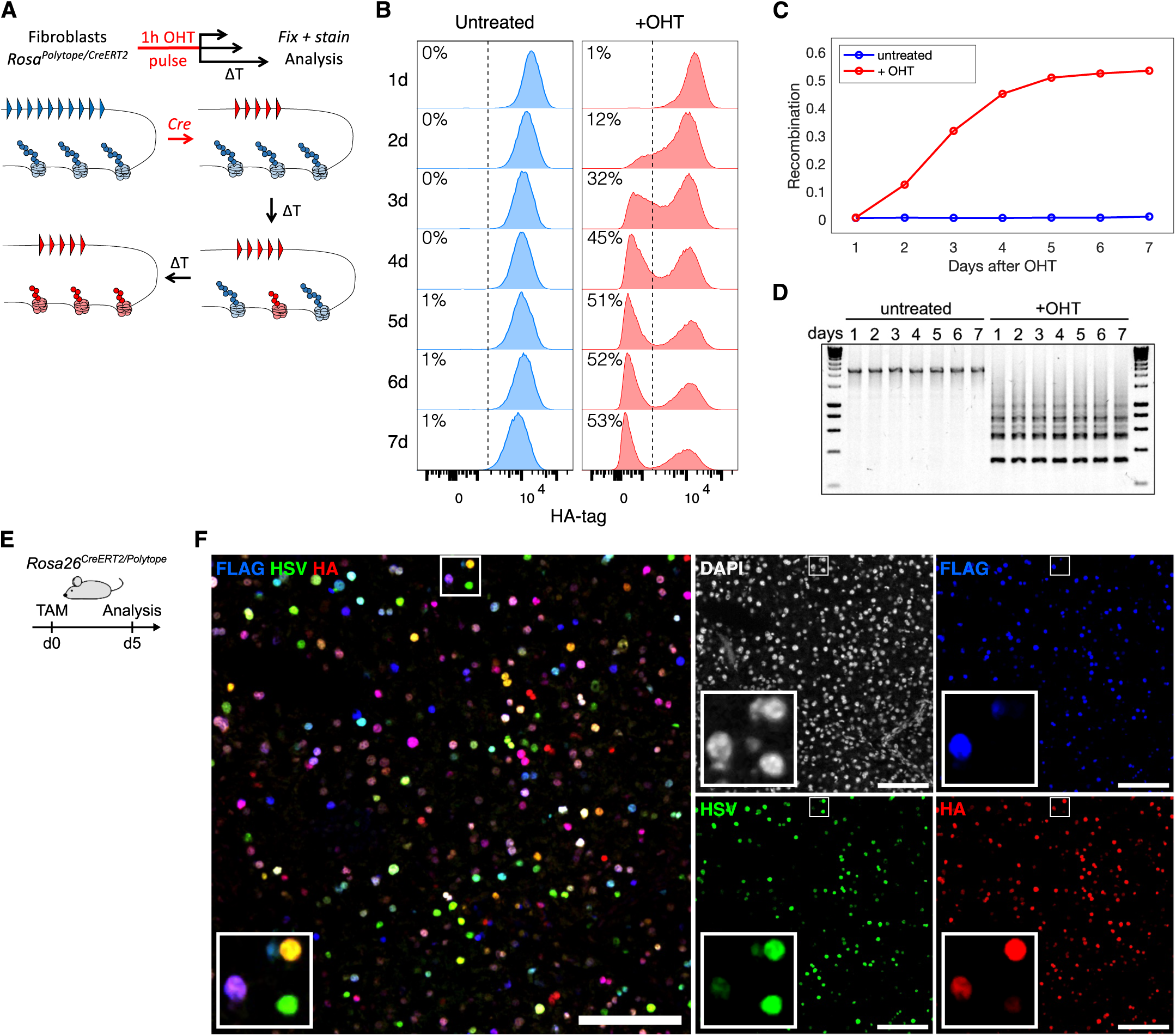
Lag time between completion of Cre/loxP recombination and incorporation of nascent H2B-Polytope peptides. (**A**) Experimental design (top): immortalized fibroblasts derived from *Rosa26^CreERT2/Polytope^* mice were treated for 1h with OHT *in vitro* and analyzed at different time points by fixation, immunostaining and flow cytometry. Molecular model (bottom): Starting with the unrecombined locus and full-length H2B-Polytope peptides (blue), activation of Cre recombination results in generation of a new DNA sequence (red), resulting in subsequent exchange of old H2B-Polytope peptides (blue) with nascent H2B-Polytope molecules (red). (**B**) Quantification of mean fluorescent intensity values using HA tag as a proxy for recombination. Dotted lines represent gating threshold dividing the dataset in either positive or negative. (**C**) Fraction of recombined cells (HA^neg^) over time. (**D**) PCR of the Polytope locus at different time points. **(E)** Experimental design of the *in vivo* experiment. **(F)** Fluorescent images of pancreatic tissue. Scale bar: 100 µm.

**Suppl. Fig. 8:**
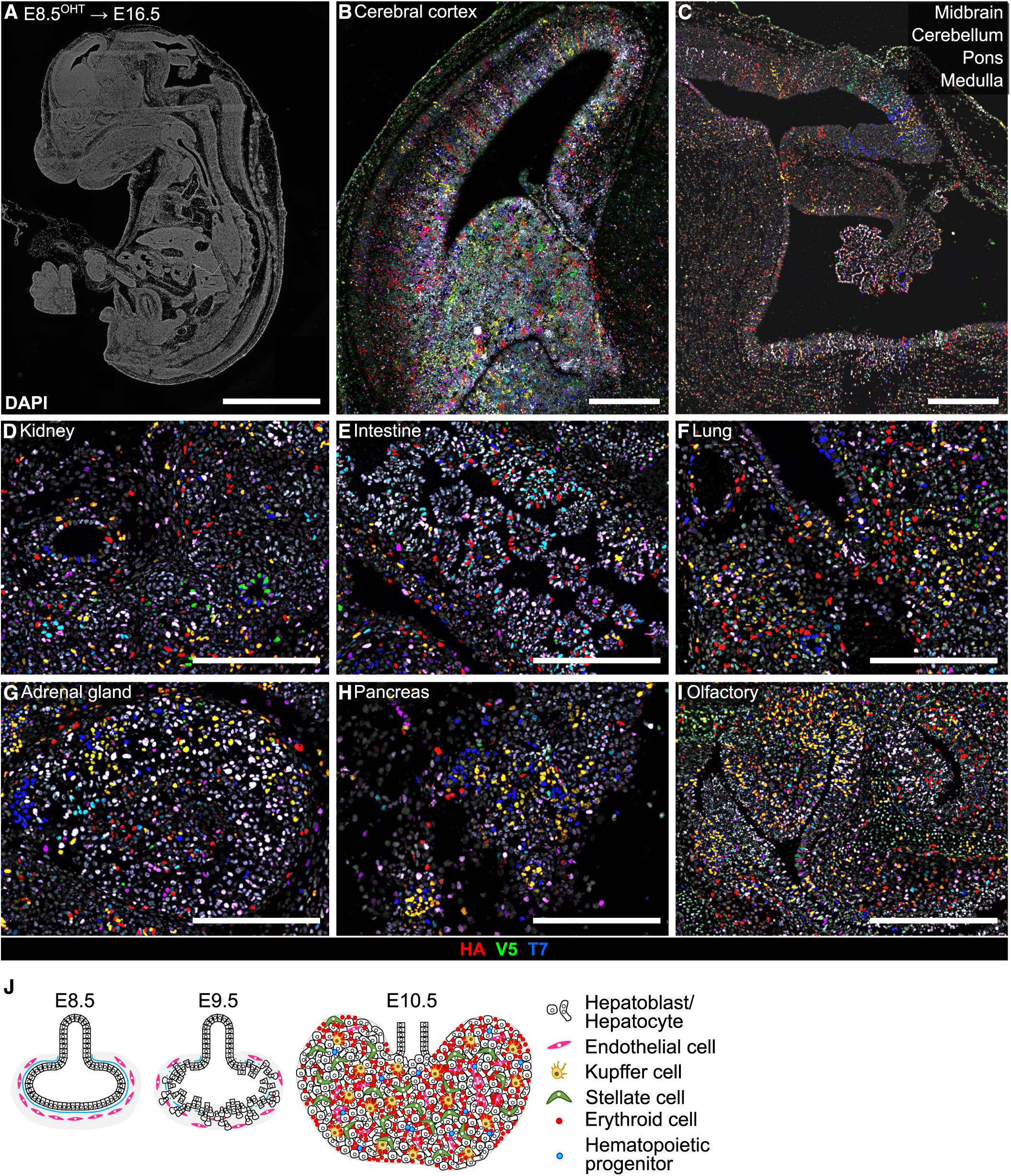
Spatial fate-mapping with Polytope in the developing embryo. *Rosa26^CreERT2/Polytope^*embryos were labeled at E8.5 and analyzed at E16.5. **(A)** Whole midline sagittal section of an embryo (DAPI staining, scale bar: 1 mm). **(B-I)** Images of Polytope color codes based on a triple immunostaining (HA/V5/T7) of the cerebral cortex, midbrain area (scale bar: 500 µm), kidney, intestine, lung, adrenal gland, pancreas and olfactory system (scale bar: 250 µm). **(J)** Schematic of early liver development.

**Suppl. Fig. 9:**
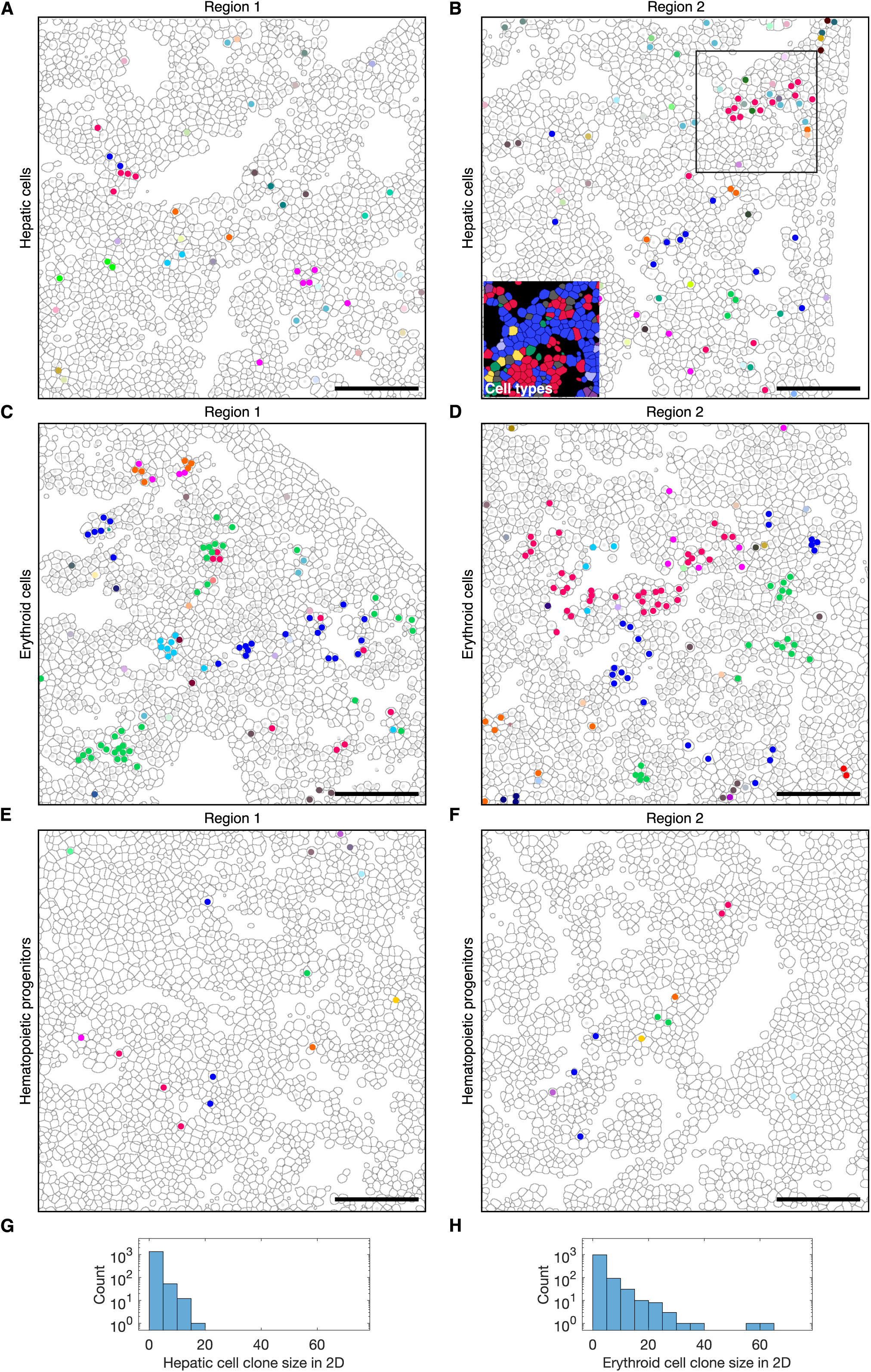
Virtual color code map filtered for specific cell types of the fetal liver barcoded at E10.5 and analyzed at E16.5. (**A-B**) Hepatic cell specific color code clusters (indicated by different colors) remapped on segmentation results. Two distinct regions depicted. Insert in (B) depicts cell type identity of cells in the box (blue: Hepatic cells; red: erythroid cells; yellow: hematopoietic progenitors; green: macrophages; purple: myeloid cells; light purple: remaining immune cells; grey: unidentified). (**C-D**) Erythroid cell specific color code clusters (indicated by different colors) remapped on segmentation results. Two distinct regions depicted. (**E-F**) Hematopoietic progenitor specific color code clusters (indicated by different colors) remapped on segmentation results. Two distinct regions depicted. Scale bar: 100 µm. **(G)** Clone size distribution in 2D for hepatic cells and **(H)** erythroid cells.

**Suppl. Fig. 10:**
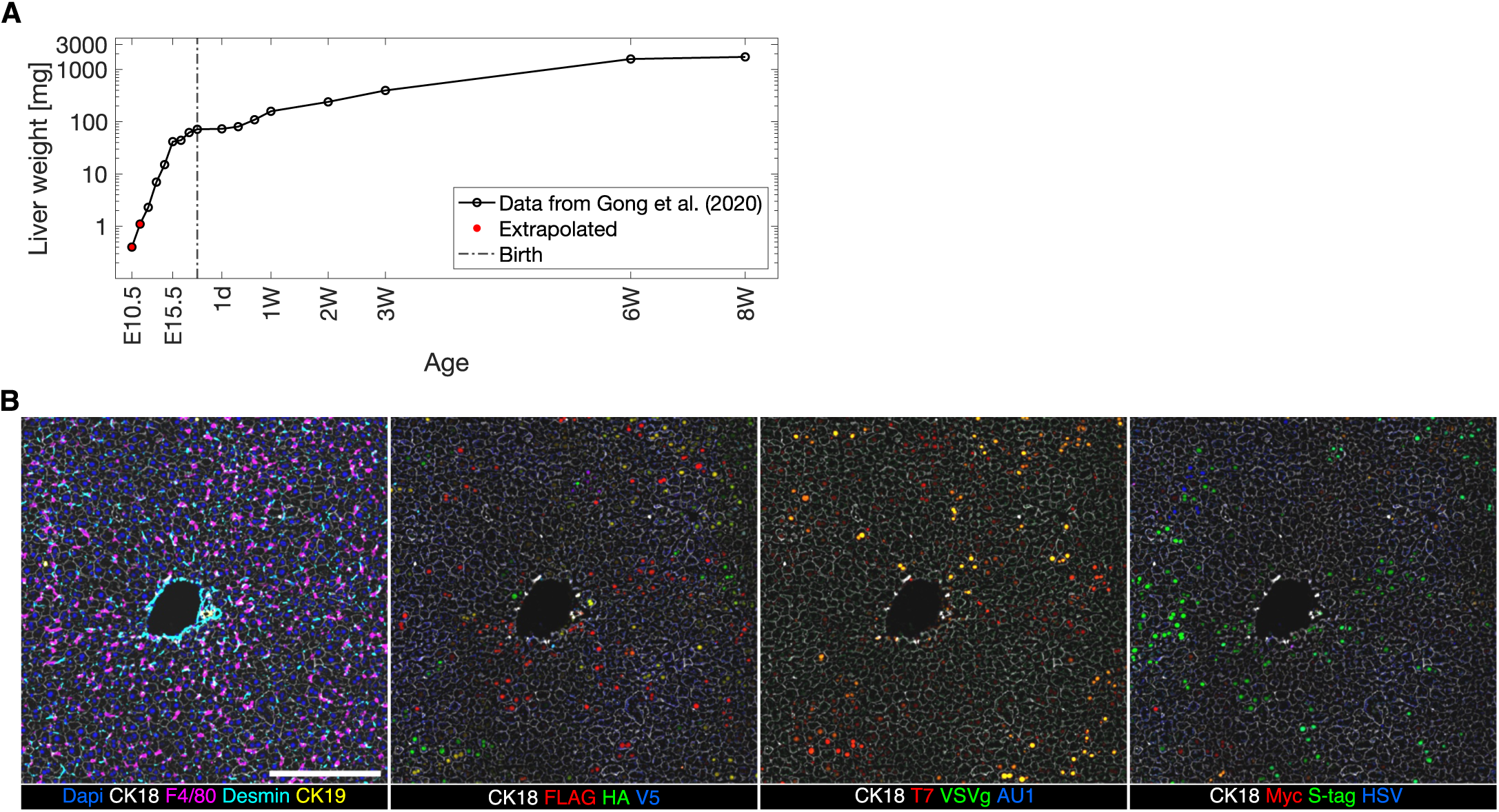
Liver development and Polytope color coding. **(A)** Liver weight during development. Based on (*42*). (**A**) Representative multiplexed images of cell marker and Polytope epitope immunostainings. Scale bar: 250 µm

**Suppl. Fig. 11:**
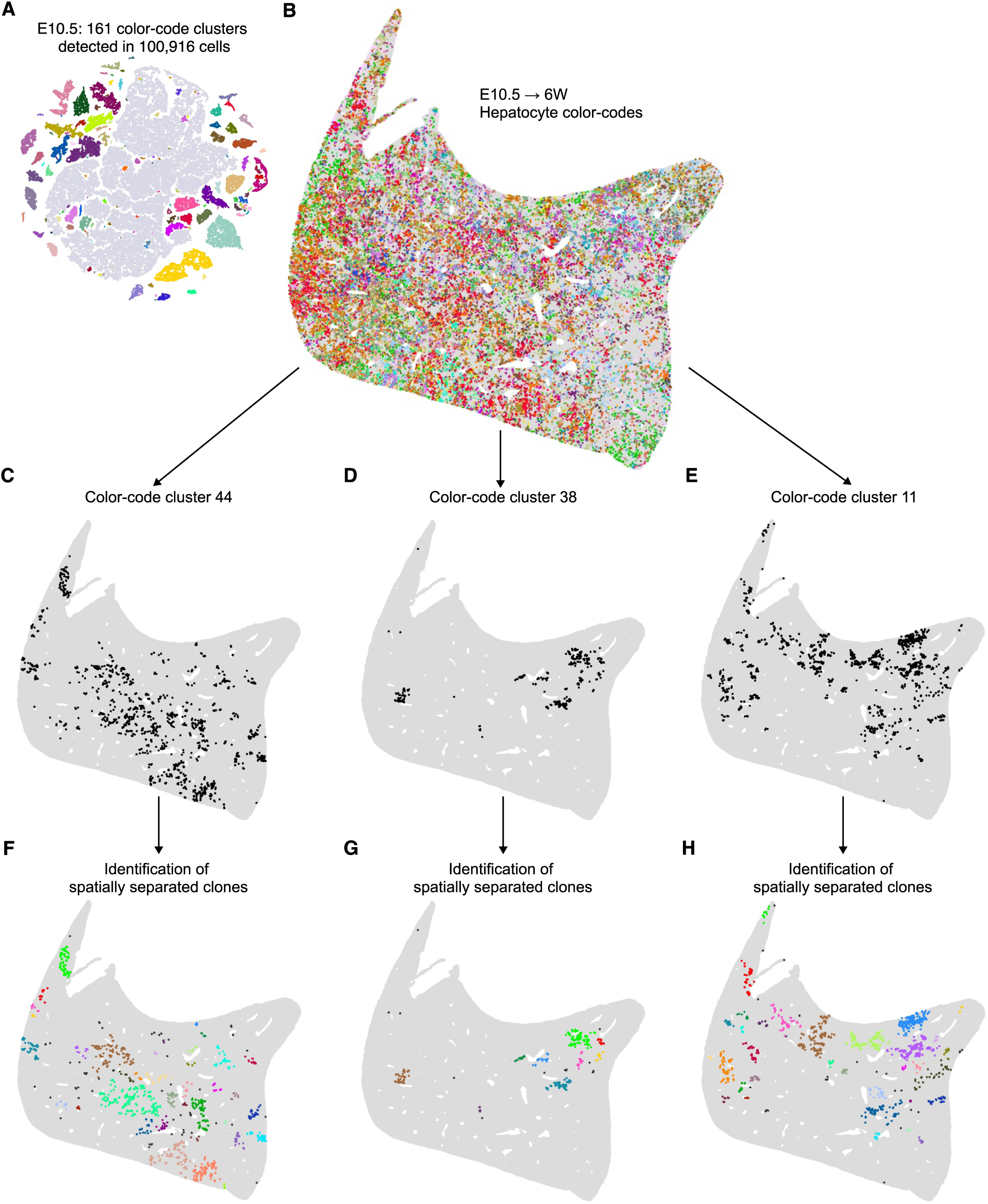
Spatial fate-mapping of E10.5 hepatocyte progenitors until adulthood. (**A**) t-SNE plot of normalized mean fluorescent intensity value and cluster detection via DBSCAN. Different colors indicate distinct color code clusters. (**B**) Remapped result from (A). (**C-E**) Remapped results of individual color code clusters and (**F-H**) detection of spatially separated clonal patches.

**Suppl. Fig. 12:**
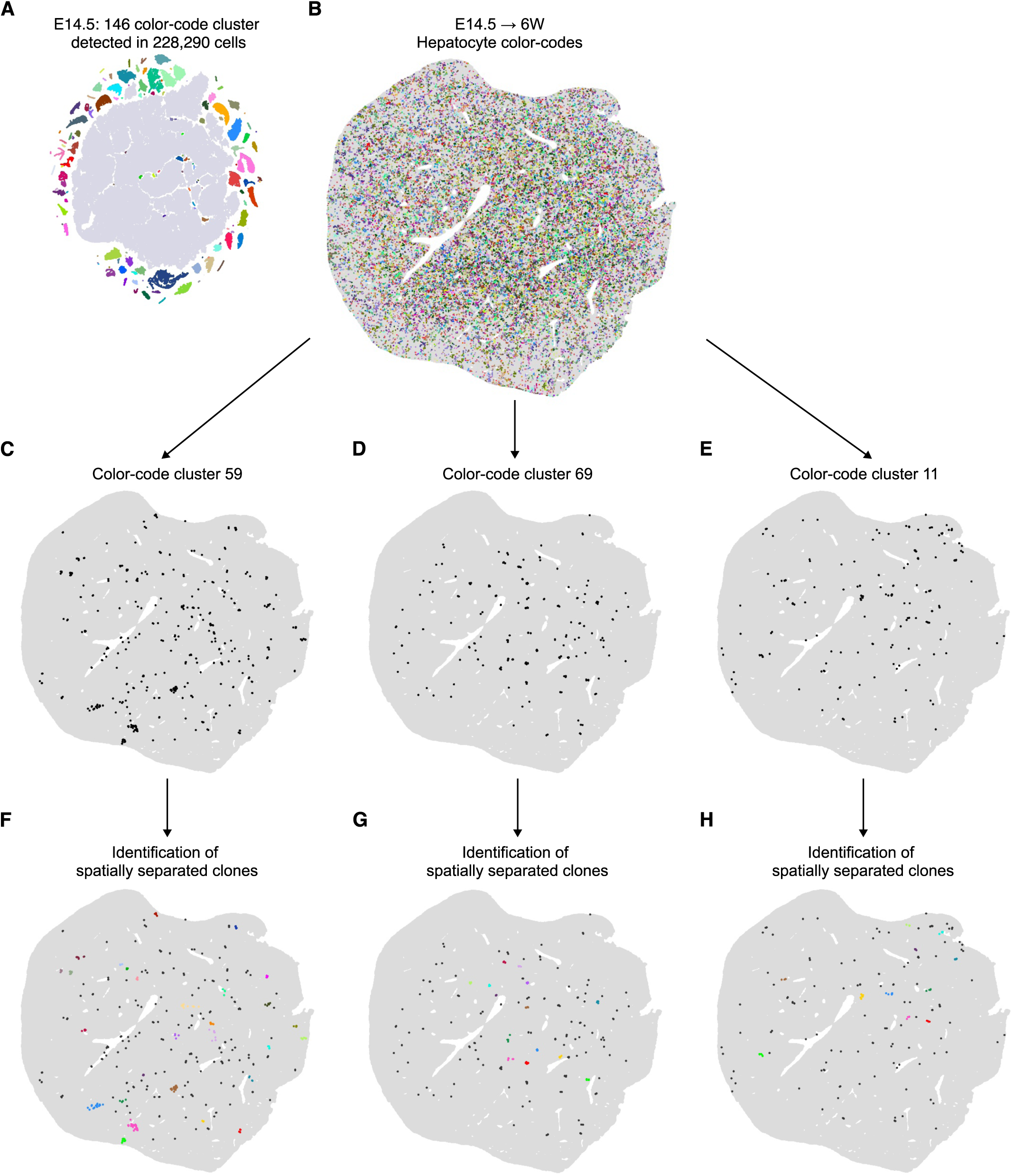
Spatial fate-mapping of E14.5 hepatocyte progenitors until adulthood. (**A**) t-SNE plot of normalized mean fluorescent intensity value and cluster detection via DBSCAN. Different colors indicate distinct color code clusters. (**B**) Remapped result from (A). (**C-E**) Remapped results of individual color code clusters and (**F-H**) detection of spatially separated clonal patches.

**Suppl. Fig. 13:**
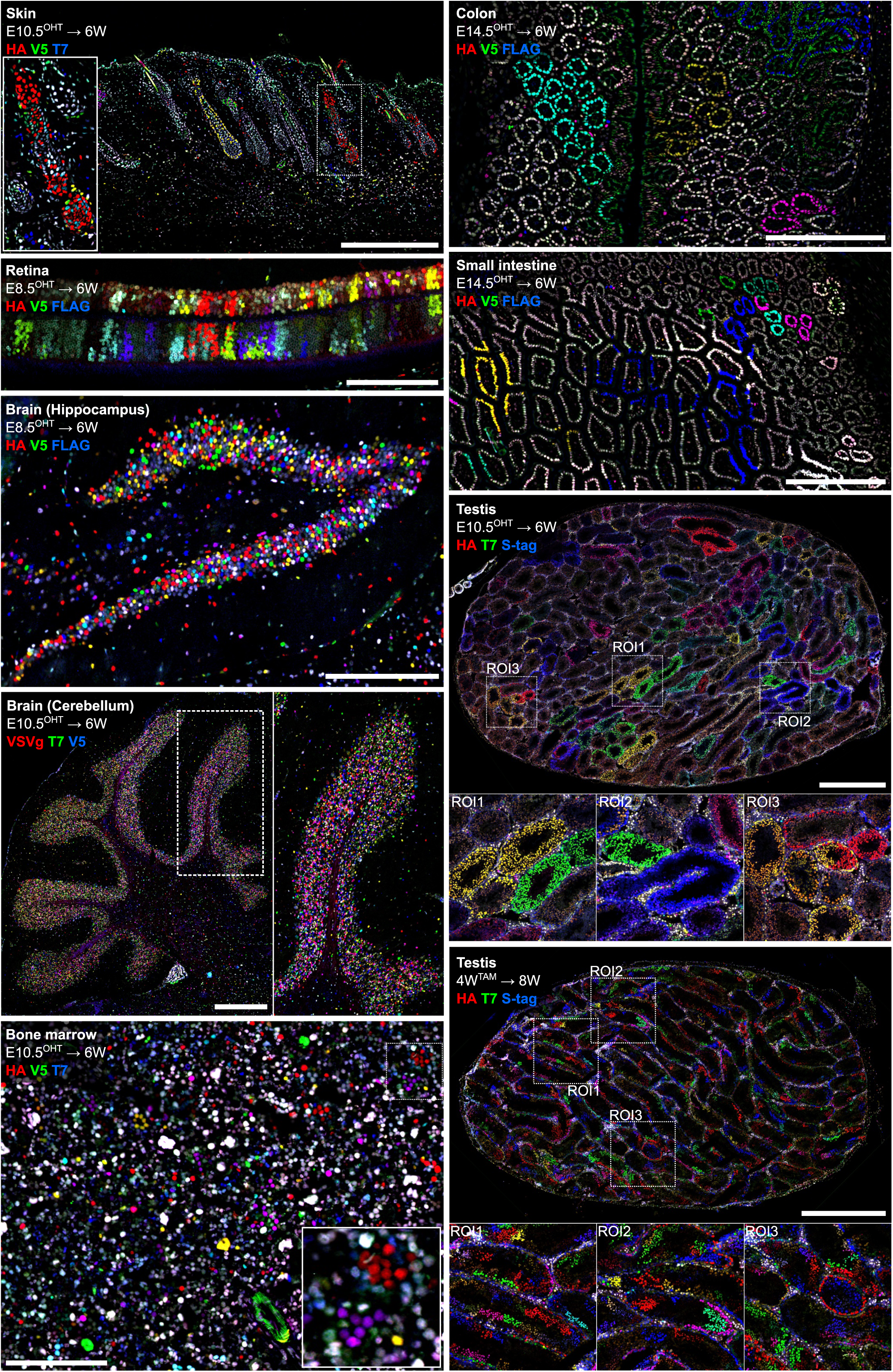
Representative images of Polytope spatial fate-mapping of embryonic progenitors until adulthood. Image collection of samples derived from *in utero* or postnatally barcoded *Rosa26^CreERT2/Polytope^*mice. Scale bars: skin, 500 µm; retina, 500 µm; hippocampus, 250 µm; cerebellum, 500 µm; bone marrow, 250 µm; colon, 250 µm; small intestine, 250 µm; testis, 1 mm.

## Notes

### Competing Interest Statement

The authors have declared no competing interest.

## References

1. T. Stadler, O. G. Pybus, M. P. H. Stumpf, Phylodynamics for cell biologists. Science (1979) 371 (2021).

2. S. Domcke, J. Shendure, A reference cell tree will serve science better than a reference cell atlas. Cell 186, 1103–1114 (2023).

3. S. VanHorn, S. A. Morris, Next-Generation Lineage Tracing and Fate Mapping to Interrogate Development. Dev Cell 56, 7–21 (2021).

4. S. E. Lee, B. D. Rudd, N. L. Smith, Fate-mapping mice: new tools and technology for immune discovery. Trends Immunol 43, 195–209 (2022).

5. J. Garcia-Marques, I. Espinosa-Medina, T. Lee, The art of lineage tracing: From worm to human. Prog Neurobiol 199, 101966 (2021).

6. N. Barker, J. H. Van Es, J. Kuipers, P. Kujala, M. Van Den Born, M. Cozijnsen, A. Haegebarth, J. Korving, H. Begthel, P. J. Peters, H. Clevers, Identification of stem cells in small intestine and colon by marker gene Lgr5. Nature 449, 1003–1007 (2007).

7. K. Busch, K. Klapproth, M. Barile, M. Flossdorf, T. Holland-Letz, S. M. Schlenner, M. Reth, T. Höfer, H. R. Rodewald, Fundamental properties of unperturbed haematopoiesis from stem cells in vivo. Nature 518, 542–546 (2015).

8. J. Livet, T. A. Weissman, H. Kang, R. W. Draft, J. Lu, R. A. Bennis, J. R. Sanes, J. W. Lichtman, Transgenic strategies for combinatorial expression of fluorescent proteins in the nervous system. Nature 450, 56–62 (2007).

9. H. J. Snippert, L. G. van der Flier, T. Sato, J. H. van Es, M. van den Born, C. Kroon-Veenboer, N. Barker, A. M. Klein, J. van Rheenen, B. D. Simons, H. Clevers, Intestinal crypt homeostasis results from neutral competition between symmetrically dividing Lgr5 stem cells. Cell 143, 134–144 (2010).

10. W. Pei, T. B. Feyerabend, J. Rössler, X. Wang, D. Postrach, K. Busch, I. Rode, K. Klapproth, N. Dietlein, C. Quedenau, W. Chen, S. Sauer, S. Wolf, T. Höfer, H. Rodewald, Polylox barcoding reveals haematopoietic stem cell fates realized in vivo. Nature 548, 456–460 (2017).

11. W. Pei, F. Shang, X. Wang, A. K. Fanti, A. Greco, K. Busch, K. Klapproth, Q. Zhang, C. Quedenau, S. Sauer, T. B. Feyerabend, T. Höfer, H. R. Rodewald, Resolving Fates and Single-Cell Transcriptomes of Hematopoietic Stem Cell Clones by PolyloxExpress Barcoding. Cell Stem Cell 27, 383–395.e8 (2020).

12. L. Li, S. Bowling, S. E. McGeary, Q. Yu, B. Lemke, K. Alcedo, Y. Jia, X. Liu, M. Ferreira, A. M. Klein, S. W. Wang, F. D. Camargo, A mouse model with high clonal barcode diversity for joint lineage, transcriptomic, and epigenomic profiling in single cells. Cell 186, 5183–5199.e22 (2023).

13. S. Bowling, D. Sritharan, F. G. Osorio, M. Nguyen, P. Cheung, A. Rodriguez-Fraticelli, S. Patel, W. C. Yuan, Y. Fujiwara, B. E. Li, S. H. Orkin, S. Hormoz, F. D. Camargo, An Engineered CRISPR-Cas9 Mouse Line for Simultaneous Readout of Lineage Histories and Gene Expression Profiles in Single Cells. Cell 181, 1410–1422.e27 (2020).

14. T. S. Weber, C. Biben, D. C. Miles, S. Glaser, S. Tomei, S. Zhang, P. P. L Tam, S. H. Naik, LoxCode in vivo barcoding resolves epiblast clonal fate to fetal organs. bioRxiv, doi: 10.1101/2023.01.02.522501 (2023).

15. B. Spanjaard, B. Hu, N. Mitic, P. Olivares-Chauvet, S. Janjuha, N. Ninov, J. P. Junker, Simultaneous lineage tracing and cell-type identification using CrIsPr-Cas9-induced genetic scars. Nat Biotechnol 36, 469–473 (2018).

16. A. McKenna, G. M. Findlay, J. A. Gagnon, M. S. Horwitz, A. F. Schier, J. Shendure, Whole-organism lineage tracing by combinatorial and cumulative genome editing. Science (1979) 353 (2016).

17. B. Raj, D. E. Wagner, A. McKenna, S. Pandey, A. M. Klein, J. Shendure, J. A. Gagnon, A. F. Schier, Simultaneous single-cell profiling of lineages and cell types in the vertebrate brain. Nat Biotechnol 36, 442–450 (2018).

18. M. M. Chan, Z. D. Smith, S. Grosswendt, H. Kretzmer, T. M. Norman, B. Adamson, M. Jost, J. J. Quinn, D. Yang, M. G. Jones, A. Khodaverdian, N. Yosef, A. Meissner, J. S. Weissman, Molecular recording of mammalian embryogenesis. Nature 570, 77–82 (2019).

19. D. E. Wagner, C. Weinreb, Z. M. Collins, J. A. Briggs, S. G. Megason, A. M. Klein, Single-cell mapping of gene expression landscapes and lineage in the zebrafish embryo. Science (1979) 360, 981–987 (2018).

20. D. Bressan, G. Battistoni, G. J. Hannon, The dawn of spatial omics. Science (1979) 381 (2023).

21. F. S. Dezem, W. Arjumand, H. DuBose, N. S. Morosini, J. Plummer, Spatially Resolved Single-Cell Omics: Methods, Challenges, and Future Perspectives. Annu Rev Biomed Data Sci 7, 131–153 (2024).

22. X. Chen, J. L. Zaro, W. C. Shen, Fusion protein linkers: Property, design and functionality. Adv Drug Deliv Rev 65, 1357–1369 (2013).

23. G. Li, Z. Huang, C. Zhang, B. J. Dong, R. H. Guo, H. W. Yue, L. T. Yan, X. H. Xing, Construction of a linker library with widely controllable flexibility for fusion protein design. Appl Microbiol Biotechnol 100, 215–225 (2016).

24. N. Sternberg, D. Hamilton, Bacteriophage P1 site-specific recombination. I. Recombination between loxP sites. J Mol Biol 150, 467–486 (1981).

25. J. R. Smith, S. Maguire, L. A. Davis, M. Alexander, F. Yang, S. Chandran, C. ffrench-Constant, R. A. Pedersen, Robust, Persistent Transgene Expression in Human Embryonic Stem Cells Is Achieved with AAVS1-Targeted Integration. Stem Cells 26, 496–504 (2008).

26. A. J. Radtke, C. J. Chu, Z. Yaniv, L. Yao, J. Marr, R. T. Beuschel, H. Ichise, A. Gola, J. Kabat, B. Lowekamp, E. Speranza, J. Croteau, N. Thakur, D. Jonigk, J. L. Davis, J. M. Hernandez, R. N. Germain, IBEX: an iterative immunolabeling and chemical bleaching method for high-content imaging of diverse tissues. Nat Protoc 17, 378–401 (2022).

27. A. J. Radtke, E. Kandov, B. Lowekamp, E. Speranza, C. J. Chu, A. Gola, N. Thakur, R. Shih, L. Yao, Z. R. Yaniv, R. T. Beuschel, J. Kabat, J. Croteau, J. Davis, J. M. Hernandez, R. N. Germain, IBEX: A versatile multiplex optical imaging approach for deep phenotyping and spatial analysis of cells in complex tissues. Proc Natl Acad Sci U S A 117, 33455–33465 (2020).

28. M. Peitz, K. Pfannkuche, K. Rajewsky, F. Edenhofer, Ability of the hydrophobic FGF and basic TAT peptides to promote cellular uptake of recombinant Cre recombinase: A tool for efficient genetic engineering of mammalian genomes. Proc Natl Acad Sci U S A 99, 4489–4494 (2002).

29. A. Foudi, K. Hochedlinger, D. Van Buren, J. W. Schindler, R. Jaenisch, V. Carey, H. Hock, Analysis of histone 2B-GFP retention reveals slowly cycling hematopoietic stem cells. Nat Biotechnol 27, 84–90 (2009).

30. Y. Li, S. Ai, X. Yu, C. Li, X. Li, Y. Yue, Y. Wei, C. Y. Li, A. He, Replication-independent histone turnover underlines the epigenetic homeostasis in adult heart. Circ Res 125, 198–208 (2019).

31. J. Lotto, T. L. Stephan, P. A. Hoodless, Fetal liver development and implications for liver disease pathogenesis. Nat Rev Gastroenterol Hepatol 20, 561–581 (2023).

32. L. Wilding Crawford, J. F. Foley, S. A. Elmore, Histology atlas of the developing mouse hepatobiliary system with emphasis on embryonic days 9.5-18.5. Toxicol Pathol 38, 872–906 (2010).

33. T. L. Tay, D. Mai, J. Dautzenberg, F. Fernández-Klett, G. Lin, S. Sagar, M. Datta, A. Drougard, T. Stempfl, A. Ardura-Fabregat, O. Staszewski, A. Margineanu, A. Sporbert, L. M. Steinmetz, J. A. Pospisilik, S. Jung, J. Priller, D. Grün, O. Ronneberger, M. Prinz, A new fate mapping system reveals context-dependent random or clonal expansion of microglia. Nat Neurosci 20, 793–803 (2017).

34. J. Zhang, Q. Wu, C. B. Johnson, G. Pham, J. M. Kinder, A. Olsson, A. Slaughter, M. May, B. Weinhaus, A. D’Alessandro, J. D. Engel, J. X. Jiang, J. M. Kofron, L. F. Huang, V. B. S. Prasath, S. S. Way, N. Salomonis, H. L. Grimes, D. Lucas, In situ mapping identifies distinct vascular niches for myelopoiesis. Nature 590, 457–462 (2021).

35. K. Khodosevich, J. Alfonso, H. Monyer, Dynamic changes in the transcriptional profile of subventricular zone-derived postnatally born neuroblasts. Mech Dev 130, 424–432 (2013).

36. L. Frank, D. Postrach, M. Langhinrichs, T. Höfer, H.-R. Rodewald, Holistic cellular barcoding reveals a common lineage tree of tissue macrophage development. bioRxiv (in preparation) (2024).

37. H. Luche, O. Weber, T. N. Rao, C. Blum, H. J. Fehling, Faithful activation of an extra-bright red fluorescent protein in “knock-in” Cre-reporter mice ideally suited for lineage tracing studies. Eur J Immunol 37, 43–53 (2007).

38. S. Ma, J. E. Hernandez, W. J. M. Huang, Protocol to assess cell-intrinsic regulatory mechanisms using an ex vivo murine T cell polarization and co-culture system. STAR Protoc 3 (2022).

39. J. Schindelin, I. Arganda-Carreras, E. Frise, V. Kaynig, M. Longair, T. Pietzsch, S. Preibisch, C. Rueden, S. Saalfeld, B. Schmid, J. Y. Tinevez, D. J. White, V. Hartenstein, K. Eliceiri, P. Tomancak, A. Cardona, Fiji: An open-source platform for biological-image analysis. Nat Methods 9, 676–682 (2012).

40. C. Stringer, T. Wang, M. Michaelos, M. Pachitariu, Cellpose: a generalist algorithm for cellular segmentation. Nat Methods 18, 100–106 (2021).

41. N. Rahmah, I. S. Sitanggang, “Determination of Optimal Epsilon (Eps) Value on DBSCAN Algorithm to Clustering Data on Peatland Hotspots in Sumatra” in IOP Conference Series: Earth and Environmental Science (Institute of Physics Publishing, 2016)vol. 31.

42. T. Gong, C. Zhang, X. Ni, X. Li, J. Li, M. Liu, D. Zhan, X. Xia, L. Song, Q. Zhou, C. Ding, J. Qin, Y. Wang, A time-resolved multi-omic atlas of the developing mouse liver. Genome Res 30, 263–275 (2020).

